# Structural insights into the human P2X1 receptor and ligand interactions

**DOI:** 10.1101/2024.04.04.588192

**Authors:** Felix M. Bennetts, Hariprasad Venugopal, Alisa Glukhova, Jesse I. Mobbs, Sabatino Ventura, David M. Thal

## Abstract

The P2X1 receptor is a trimeric ligand-gated ion channel that plays a pivotal role in urogenital and immune functions. Consequently, it offers numerous potential indications for novel drug treatments. Unfortunately, the progress of drug discovery targeting the P2X1 receptor has been impeded by the absence of structural information. To gain deeper insights into the binding site of the P2X1 receptor, we employed cryogenic electron microscopy (cryo-EM) to elucidate the structures of the P2X1 receptor in both an ATP-bound desensitised state and an NF449-bound closed state. NF449 is a potent P2X1 receptor antagonist and engages with the receptor in a distinctive manner. To gain insights into the molecular machinery governing receptor inhibition and activation and better understand P2X1 receptor ligand subtype selectivity, critical P2X1 receptor residues involved in ligand binding were mutated. Radioligand binding assays with [^3^H]-α,β-methylene ATP and intracellular calcium influx were employed to assess the effect of these mutations on ligand binding and receptor activation, thereby validating key ligand-receptor interactions. This research expands our understanding of the P2X1 receptor structure at a molecular level and opens new avenues for *in silico* drug design targeting the P2X1 receptor.

## Introduction

There are seven subtypes of the P2X ligand-gated ion channel (P2X1-7) and these receptors control a vast number of biological processes that make them exciting drug targets.^1^ P2X receptors consist of three individual subunits that can be homotrimeric or heterotrimeric to form a ligand-gated ion channel.^2,3^ Each P2X subunit contains two transmembrane-spanning helices, a large extracellular domain, and a small intracellular N- and C-terminus, except for the P2X7 receptor, which has a longer C-terminus. The endogenous ligand, ATP, activates P2X receptors by binding to ATP binding sites located at the extracellular interface of each receptor subunit. Upon activation, P2X receptors form a non-selective pore that is permeable to cations such as potassium, calcium, and sodium.^4^

Research among the P2X receptor family has primarily focused on the P2X3, P2X4 and P2X7 receptors.^5^ Experimentally determined structures exist for each of these P2X receptor subtypes alongside a selection of potent and specific small molecule compounds.^5–11^ These groundbreaking studies have greatly improved our understanding of the P2X receptors. However, for some of the lesser studied P2X receptors, crucial information is still lacking, particularly regarding molecular mechanisms of activation and deactivation and the relationship between structure and function, with a focus on how this knowledge could be utilised for drug discovery.^12^ The P2X1 receptor is among these receptors, known for its high affinity for the endogenous ligand ATP and its rapid desensitisation following receptor activation.^5^ The P2X1 receptor holds therapeutic promise, particularly in the urogenital and immune system. P2X1 receptor knockouts have revealed reduced fertility in male mice, attributed to diminished contractility of the vas deferens, suggesting a P2X1 receptor antagonist could be used as a male contraceptive.^13,14^ The P2X1 receptor’s involvement in platelets and neutrophils contributes to thrombo-inflammatory conditions, which could potentially be targeted by a drug to suppress clot formation, thereby serving as an antithrombotic agent.^15,16^ Recent studies have even indicated the involvement of P2X1 receptors in different types of cancers and small arteries, hinting at potential medical applications yet to be discovered.^17–19^

In recent decades, research groups have employed structure-activity relationship studies with known P2X1 receptor ligands or screening methods to discover improved P2X1 receptor ligands. The most promising candidates include PSB-2001, ATA, MRS2159, and the compound series introduced by Jung et al.^20–23^ Yet, none are as potent or P2X receptor selective as an older generation P2X1 receptor antagonist, NF449.^24,25^ It is important to note that NF449 also exhibits several off-target effects and is a large polar molecule making it difficult to improve.^26^ Nevertheless, NF449 is an interesting molecule that may be useful in understanding how P2X1 receptor antagonists can mediate the inhibition of the receptor. The majority of drug discovery efforts have been directed towards the generation of P2X1 receptor antagonists, and there are currently very few druglike P2X1 receptor agonists, suggesting potential for substantial improvement in this area.^12^ The endogenous ligand for P2X receptors is ATP and P2X receptor structures and mutagenesis studies have highlighted the essential interactions ATP forms with P2X receptors.^9–11,27^ Additionally, mutagenesis studies indicate the involvement of similar residues in ATP-mediated receptor activation at the P2X1 receptor.^27^ To explore the molecular components of activation and inhibition we have successfully determined the P2X1 receptor structure in an ATP-bound desensitised state and a NF449-bound closed state. These structures have guided mutagenesis studies aimed at unravelling ligand binding at a molecular level at the P2X1 receptor. Through comparisons with previously determined P2X receptor structures and mutagenesis, we delineate the structural features and mechanistic basis for subtype-specific interactions of the P2X1 receptor.

## Results

### Structure determination of the P2X1 receptor

The full-length human P2X1 receptor was extracted from insect cells and reconstituted into detergent micelles for single-particle cryogenic electron microscopy (Supplementary Fig. 1). Cryo-EM imaging of the P2X1 receptor revealed severe preferred orientation of the receptor in vitreous ice. The addition of fluorinated Fos-choline-8 before vitrification alleviated orientation bias and allowed for the successful determination of the P2X1 receptor to high resolution (Supplementary Fig. 2). In an early cryo-EM dataset of the P2X1 receptor, cryo-EM density was observed in the orthosteric pocket, corresponding to ATP, even though ATP was not added during the purification of the receptor. This finding mirrors observations from the initial crystal structures of the hP2X3 receptor.^11^ In a subsequent sample, ATP and Mg^2+^ were added at high concentrations to determine the ATP-bound P2X1 receptor structure (PDB: 9B73) to a resolution of 1.96 Å (Fig. 1A, 2A). The addition of the high affinity and competitive P2X1 receptor antagonist NF449^24^ was used to remove ATP-bound to detergent solubilised P2X1 receptor (Supplementary Fig. 1) and allowed for determination of the NF449-bound P2X1 receptor structure (Supplementary Fig. 2). The NF449-bound P2X1 receptor structure (PDB: 9B95) was resolved to 2.61 Å resolution (Fig. 1B, 2B). The structure of P2X receptor monomers can be likened to that of a dolphin, featuring a head domain, left and right flippers, dorsal fin, upper body, lower body and a fluke (Fig. 1C).^2^ The P2X1 receptor models correlated well with their respective density maps and assignment of backbone, side chains, ligands, metals, glycosylation and waters were possible at the achieved resolution (Supplementary Fig. 3).

**Figure 1.**
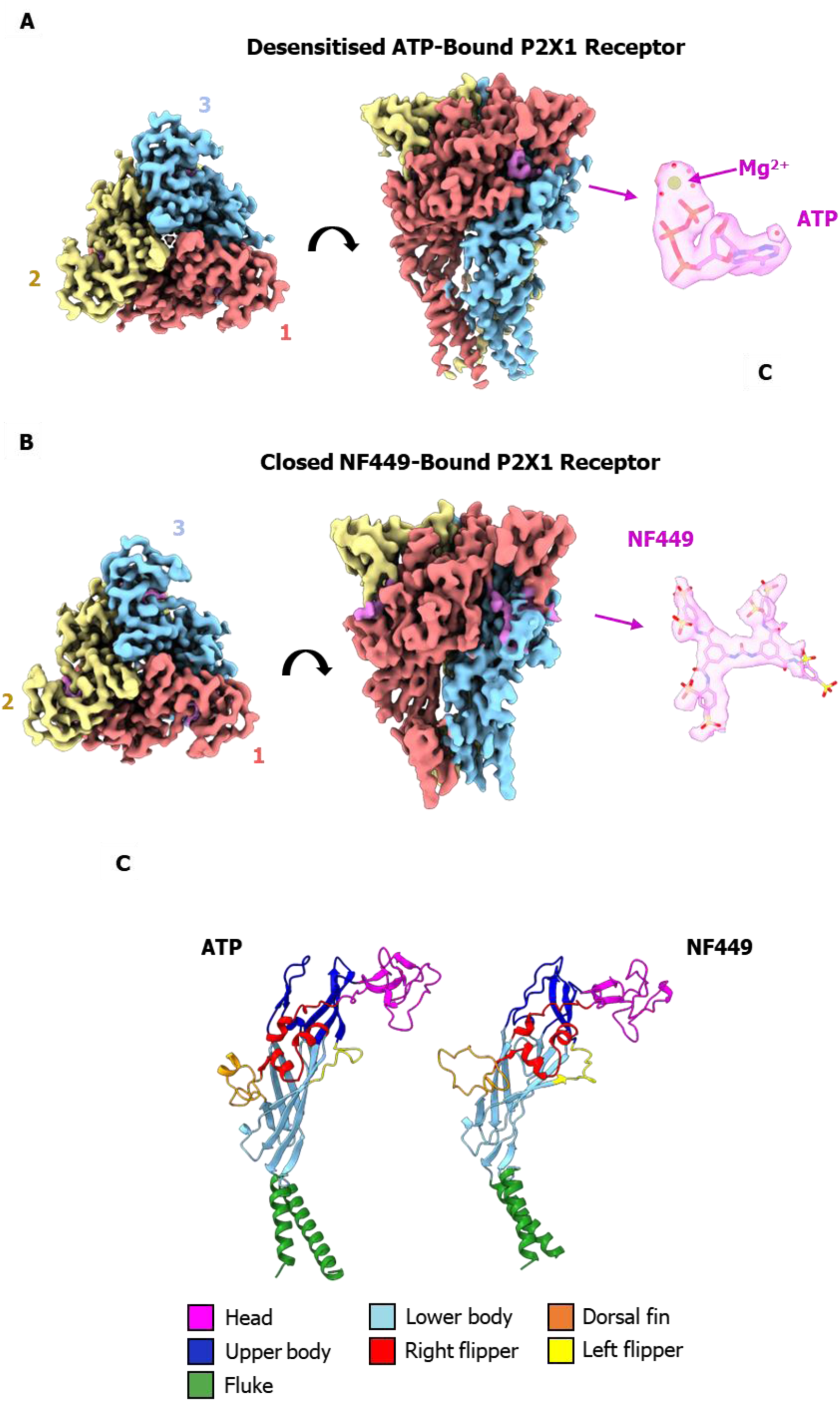
**A)** Cryo-EM map of the ATP-bound P2X1 receptor, with chains coloured individually and the cryo-EM ligand density depicted in violet. An enlarged image of Mg-ATP and nearby water molecules was fitted into the cryo-EM map. **B)** Cryo-EM map of the NF449-bound P2X1 receptor, with chains coloured individually and the cryo-EM ligand density depicted in violet. An enlarged image of NF449 was fitted into the cryo-EM map. **C)** Structural characteristics of the monomer from the ATP-bound P2X1 receptor and the NF449-bound P2X1 receptor colourised to resemble features of a dolphin. The head domain is colored purple, the lower body is light blue, the dorsal fin is orange, the upper body is blue, the right flipper is red, the left flipper is yellow, and the fluke is green.

**Figure 2.**
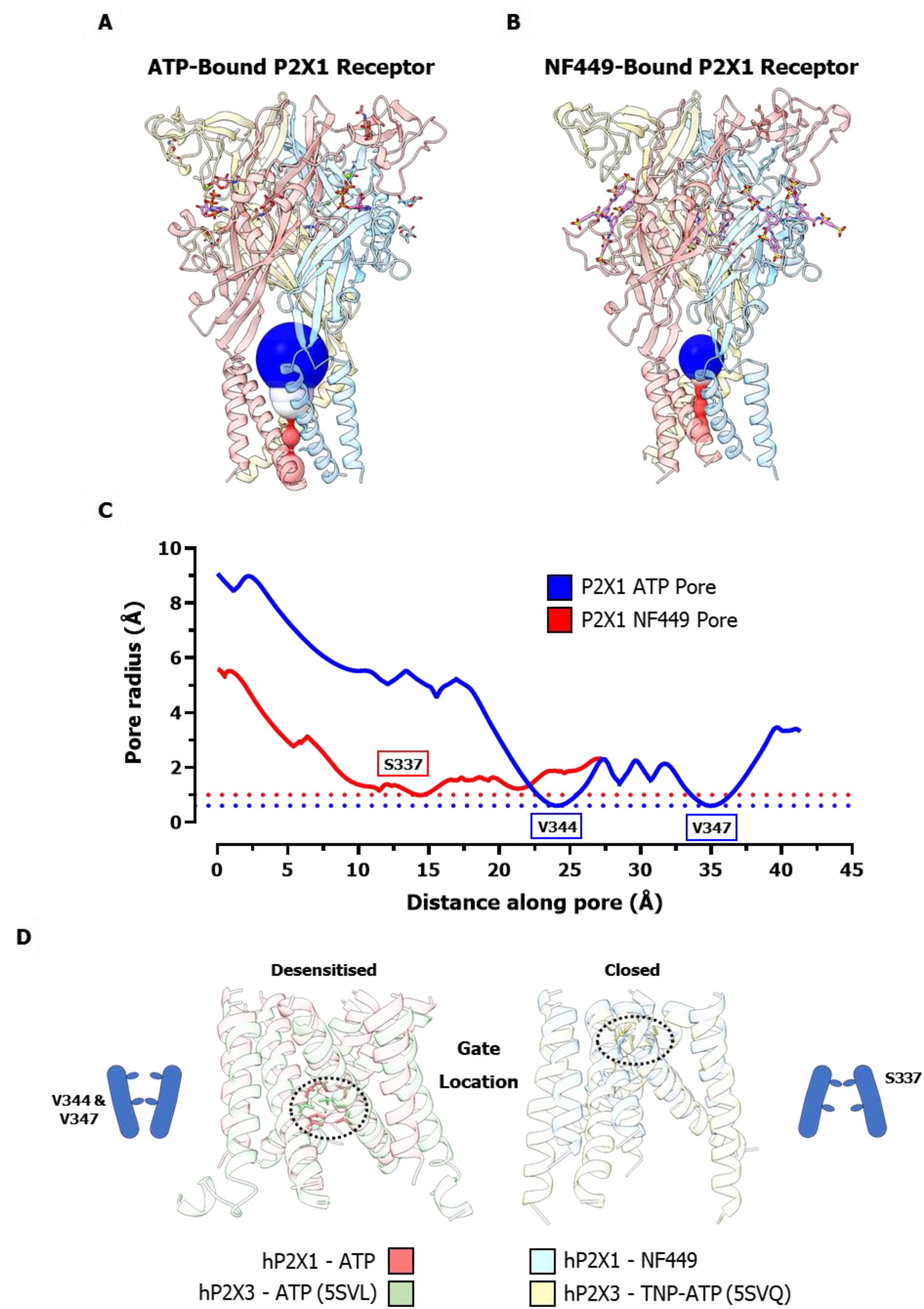
Cartoon representation of the **A)** ATP-bound P2X1 receptor and **B)** NF449-bound P2X1 receptor coloured by monomer. The surface volume of the pore radius is highlighted in red, white, and blue, with red indicating the area with the tightest constriction point. **C)** Graph of the pore radius along the length of the pore region with the residues responsible for constricting the pore radii highlighted. The ATP-bound P2X1 receptor is depicted in blue, while the NF449-bound P2X1 receptor is represented in red. The pore radius measured 0.6 Å for the ATP-bound receptor and 1.0 Å for the NF449-bound receptor, indicated by the dotted line. **D)** Transmembrane domains of the ATP-bound P2X1 receptor (red) are overlaid with those of the ATP-bound hP2X3 receptor (5SVL, green). Additionally, transmembrane domains of the NF449-bound P2X1 receptor (blue) are overlaid with those of the TNP-ATP-bound hP2X3 receptor (5SVQ, yellow).

### Structural features and gating of the P2X1 receptor

The structures of both the ATP-bound and NF449-bound P2X1 receptors exhibit a characteristic chalice-like shape, consistent with other experimentally determined P2X receptor structures (Fig. 2A, B).^9–11^ To define the state of these receptors, the pore region was analysed revealing that the ATP-bound P2X1 receptor has two gates, V344 and V347 from each monomer, which constrict the pore to its narrowest radius (Fig. 2A). Conversely, within the NF449-bound P2X1 receptor structure, S337 from each monomer constricts the pore to its narrowest radius (Fig. 2B). Both P2X1 receptor structures were in non-pore conducting states due to the small pore radius, measuring 0.6 Å for the ATP-bound receptor and 1.0 Å for the NF449-bound receptor, making it insufficient to facilitate the transport of cations (Fig. 2C). Structures of the P2X3 receptors were determined in all of the general ion channel states (desensitised, apo, open, closed).^11^ To gain deeper insights into the gating cycle of the P2X1 receptor, the desensitised and closed structures of the P2X3 receptor were compared to the P2X1 receptor structures (Fig. 2D). Similarities were noted in the location of the gate and the architecture of the transmembrane domain, suggesting a comparable state for the P2X1 receptor structures. These similarities and the pore radius indicate that the ATP-bound P2X1 receptor was in a desensitised state, while the NF449-bound P2X1 receptor was in a closed state. Another intriguing finding was the inability to model the cytoplasmic region in the P2X1 receptor structures due to a lack of cryo-EM map density, which was particularly relevant as a full-length receptor was utilised. A comparable observation was made in the structures of the P2X3 receptor in its desensitised and closed states, where the cytoplasmic region was only resolved in the open state.^11^ It is likely that this region is highly dynamic and disordered in the desensitised and closed states of the P2X1 and P2X3 receptors.

NF449 is a potent and competitive P2X1 receptor antagonist and aligning the ATP-bound P2X1 receptor and NF449-bound P2X1 receptor structures revealed one benzene disulfonic acid group of NF449 overlaping with ATP, demonstrating how NF449 competitively inhibits ATP from binding the P2X1 receptor (Fig. 3A). Initially it was difficult to visualise how a large molecule like NF449 would fit into the orthosteric binding pocket however, from the structure it was apparent that NF449 adopts a distinct orientation with unique contacts to the lower body, head domain, and dorsal fin of the P2X1 receptor (Fig. 3A). Furthermore, there were significant differences around the ligand binding sites at a structural level. The left flipper, right flipper, head domain and dorsal fin moved outward in the NF449-bound P2X1 receptor structures (Fig. 3A). The changes around the ligand binding site propagate a conformational change through the lower body to the pore region (Fig. 3B). The lower body and upper portion of the transmembrane domains moved diagonally inward and downward resulting in the vestibule above the pore reducing size and the constriction site of the pore moving up the transmembrane domain (Fig. 3B).

**Figure 3.**
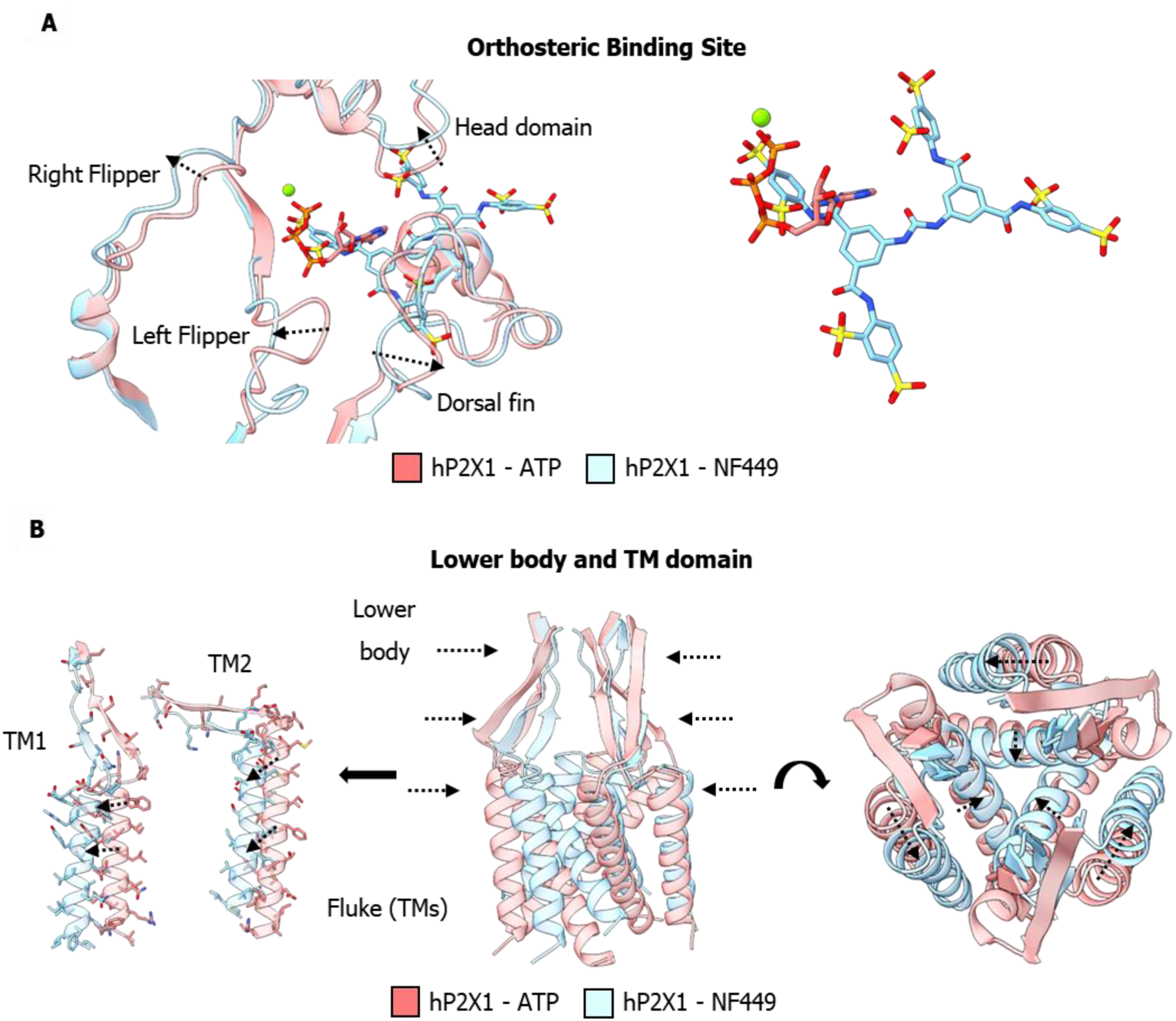
**A)** Overlay of the ATP-bound P2X1 receptor (red) and NF449-bound P2X1 receptor (blue) at the orthosteric binding site, illustrating movement in secondary structure with arrows. **B)** Overlay of the ATP-bound P2X1 receptor (red) and NF449-bound P2X1 receptor (blue) at the lower body and transmembrane domain, illustrating movement in secondary structure with arrows.

### ATP binding site of the P2X1 receptor

It is known that ATP exhibits high affinity and potency for the P2X1 receptor, which was further confirmed using a competition radioligand binding assay and a calcium influx assay using a more stable related analogue of ATP, α,β-methylene ATP. α,β-methylene ATP exhibited an affinity of 43 nM (Supplementary Fig. 5) and a potency of 99 nM (Table 1) for the P2X1 receptor. The potency of α,β-methylene ATP was comparable to previous studies^28^, while the affinity was previously reported at a lower value of 8.1 nM, with the differences likely due to different experimental conditions.^29^ The cryo-EM structure of the ATP-bound P2X1 receptor revealed a wide range of interactions between ATP and the receptor (Fig. 4A, Supplementary Table 2). Residues K68, K70, R292, and K309 formed salt bridges with the phosphate chain of ATP, with residue N290 and S286 additionally contributing a hydrogen bond in this region, collectively generating a highly polar environment around the phosphate chain of ATP. The phosphate chain of ATP formed an ionic interaction with a magnesium ion, which in turn engaged in a direct ionic interaction with residue D170, while also interacting with residue E122 through a water network, contributing to the highly polar local environment around the phosphate chain of ATP. Residues K68, K70, N290, R292, and K309 were previously studied using single alanine mutations and were shown to cause large reductions in the potency of ATP when compared to the WT-P2X1 receptor.^27^ Another critical residue was T186, which forms hydrogen bonds with the adenine ring of ATP utilising both the backbone carbonyl and the side chain hydroxyl group (Fig. 4A). Mutation of T186 was shown to be an important component of ATP-mediated receptor activation.^27^ A new finding, possible because of the high resolution of the ATP-bound P2X1 receptor structure, was a water-mediated hydrogen bond between R139 and the adenine ring of ATP. In addition, residue K140 formed a hydrogen bond with the ribose sugar of ATP, while the backbone carbonyl of K68 formed a hydrogen bond with the adenine ring of ATP. Together, residues, K68, R139, and K140 form a polar pocket around the adenine ring and ribose sugar of ATP. F188 engages in a pi-pi stacking interaction with the adenine ring, while on the opposite side of the adenine ring, K70 participates in a pi-cation interaction. Finally, hydrophobic residues, M214, and V229 form a small hydrophobic pocket around the adenine ring of ATP, with part of this region being previously studied using cysteine scanning mutagenesis and shown to be important in ATP-mediated P2X1 receptor activation.^30^

**Figure 4.**
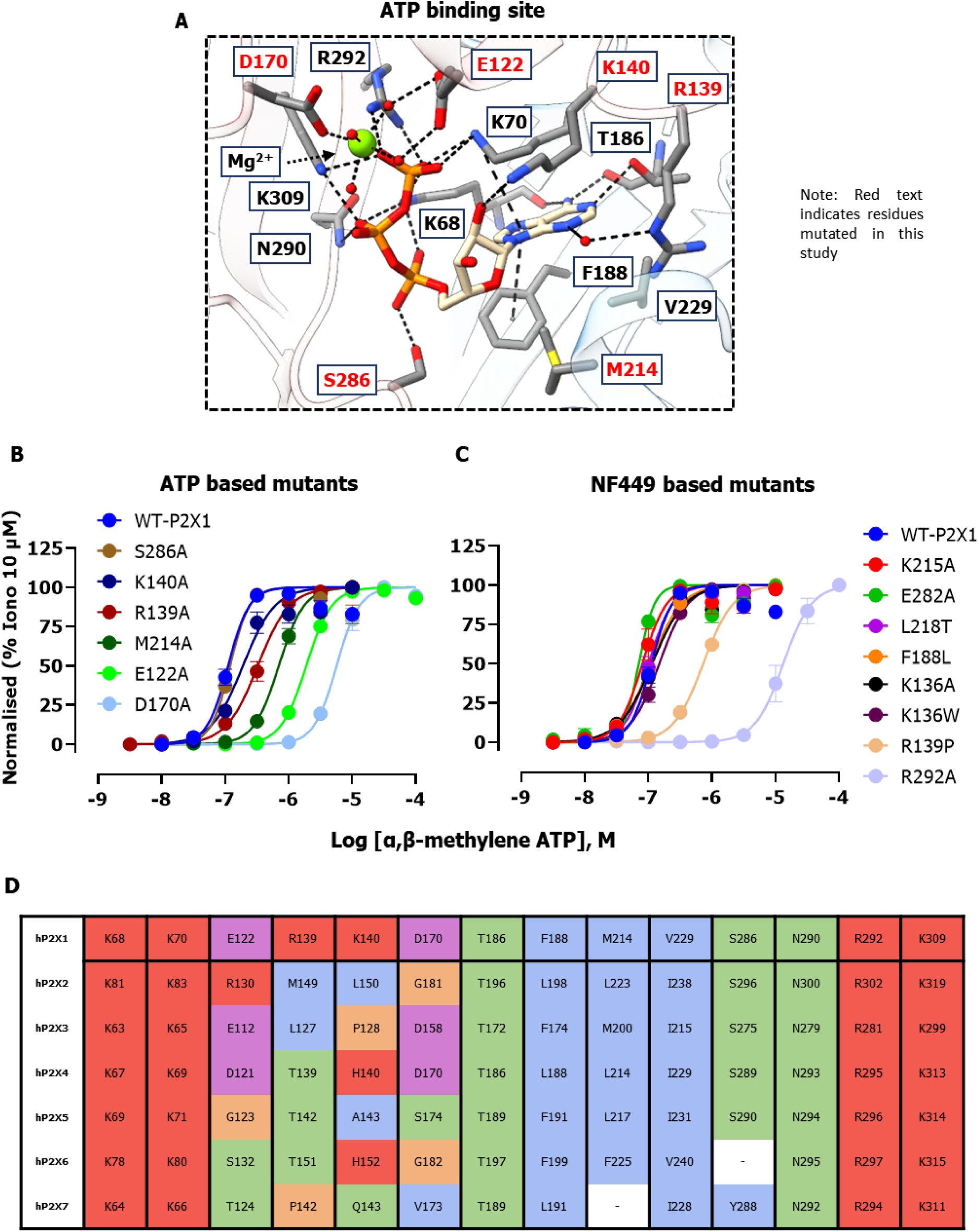
**A)** Depiction of the ATP binding site where the key interactions are illustrated as black dotted lines, accompanied by labeling of nearby residues. **B, C)** Increasing concentrations of the agonist α,β-methylene ATP on HEK293 cells expressing WT-P2X1 or single residue mutants of the P2X1 receptor. The data is normalized to 10 µM ionomycin and subsequently adjusted so that the highest value corresponds to 100 and the lowest value to 0. It is then fitted to a log(agonist) nonlinear regression curve using a four-parameter model, with the top and bottom constrained to 100 and 0, respectively. Results are presented as mean ± SEM, with n = 4 replicates. The left graph contains mutants designed for the ATP-bound P2X1 receptor and the right graph contains mutants designed for the NF449-bound P2X1 receptor. **D)** Amino acid sequence alignment of key residues from the human P2X1 receptor (hP2X1:P51575), aligned with corresponding residues from other human P2X receptor subtypes (hP2X2:Q9UBL9, hP2X3:P56373, hP2X4:Q99571, hP2X5:Q93086, hP2X6:O15547, hP2X7:Q99572). Residue categorisation based on properties: Hydrophobic (A, I, L, M, F, W, V, Y) in blue, positive charge (K, R, H) in red, negative charge (E, D) in magenta, polar (N, Q, S, T) in green. Special cases (C, G, P) in orange. Gaps are represented in white.

**Table 1.**
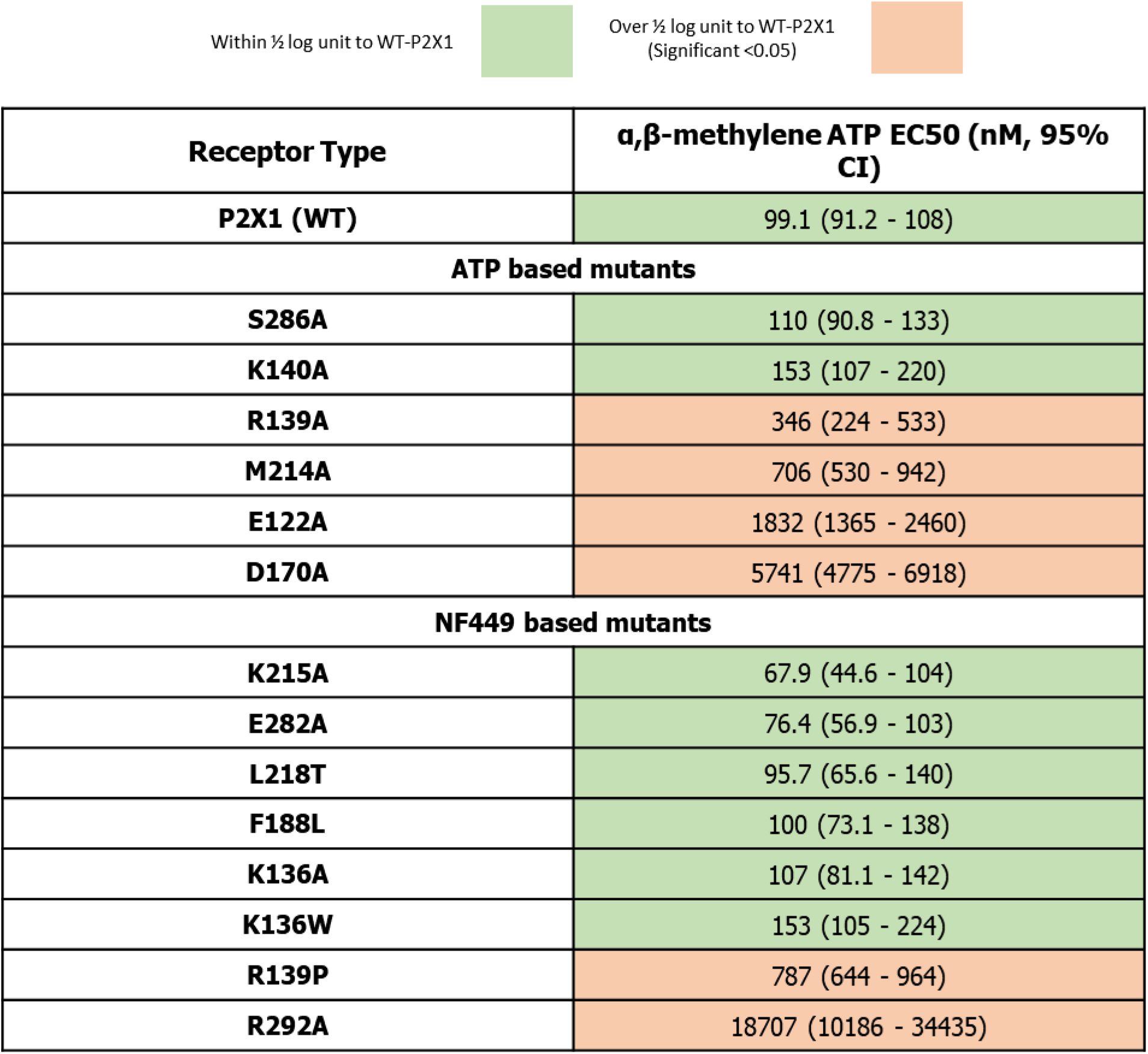
Calculated EC50 values of α,β-methylene ATP derived from pooled data of four individual experiments from HEK293 cells expressing WT-P2X1 and single residue mutants of the P2X1 receptor, alongside their respective 95% confidence intervals. Values are coloured green if within half a log unit of WT-P2X1 and coloured orange if over half a log unit of WT-P2X1 with all values over a half a log unit statistically significant as calculated by a Dunnett’s multiple comparisons test.

| Receptor Type | $\alpha,\beta$-methylene ATP EC50 (nM, 95% CI) |
| --- | --- |
| **P2X1 (WT)** | 99.1 (91.2 - 108) |
| **ATP based mutants** |  |
| **S286A** | 110 (90.8 - 133) |
| **K140A** | 153 (107 - 220) |
| **R139A** | 346 (224 - 533) |
| **M214A** | 706 (530 - 942) |
| **E122A** | 1832 (1365 - 2460) |
| **D170A** | 5741 (4775 - 6918) |
| **NF449 based mutants** |  |
| **K215A** | 67.9 (44.6 - 104) |
| **E282A** | 76.4 (56.9 - 103) |
| **L218T** | 95.7 (65.6 - 140) |
| **F188L** | 100 (73.1 - 138) |
| **K136A** | 107 (81.1 - 142) |
| **K136W** | 153 (105 - 224) |
| **R139P** | 787 (644 - 964) |
| **R292A** | 18707 (10186 - 34435) |

Although most of the ATP binding site residues were previously studied^27^ several residues, including E122, R139, K140, D170, M214, and S286 remain unexplored (Fig. 4A). Some of these residues, E122, R139, K140, and D170 show low amino acid sequence conservation across the P2X receptor family, which is uncommon in the ATP binding site (Fig 4D). To understand their role in ATP-mediated receptor activation, each of these residues was individually mutated to alanine. Subsequently, activation of the mutated receptor was assessed using calcium influx assays. The mutants R139A, M214A, E122A, D170A all significantly decreased the potency of the P2X1 agonist α,β-methylene ATP compared to WT-P2X1 with a 3.5-fold, 7-fold, 18.5-fold and 58-fold decrease respectively (Fig 4B, Table 1). In contrast, mutant S286A and K140A did not significantly change the potency of α,β-methylene ATP compared to WT-P2X1 (Fig 4B, Table 1). To validate if these mutations were driving a change in the affinity of α,β-methylene ATP WT-P2X1, each mutant receptor was tested in a radioligand binding experiment using [^3^H]-α,β-methylene ATP. The mutants S286A and K140A, which did not affect the potency of α,β-methylene ATP, exhibited binding affinity comparable to that of WT-P2X1 (Supplementary Fig. 5). Mutated residue R139A showed a 6-fold decrease in affinity for [^3^H]-α,β-methylene ATP and mutants M214A, E122A, and D170A exhibited reduced binding affinities that were beyond the threshold that could be accurately measured using a radioligand probe (Supplementary Fig. 5). These findings correlate with the substantial decrease in potency observed in the calcium activation assay for each of these mutants. This suggests that the decline in potency for the mutants R139A, M214A, E122A, and D170A stems at least in part from a decrease in the agonist’s affinity to bind to the mutated P2X1 receptor. In addition to the known residues crucial for ATP-mediated P2X1 receptor activation, residues R139, M214, E122, and D170 emerge as significant contributors to P2X1 receptor activation.^27^ Furthermore, these residues exhibit lower conservation within the P2X receptor family compared to other interactions of the ATP binding site, implying that they enhance the potency of ATP for the P2X1 receptor and could potentially be residues to target for the design of subtype-selective agonists.

To validate the presence of the magnesium ion within the ATP binding site, metal binding parameters were utilised to calculate various factors such as valency, occupancy, geometry, and distance of contacts (Supplementary Fig. 8).^31–35^ The magnesium ion was coordinated by six contacts in an octahedral geometry, four to water molecules, one to the gamma phosphate of ATP and one to D170. Each interaction formed an ionic bond at a distance of 2.0 angstroms (Supplementary Table 2). The nature and qualities of these interactions suggest this site was likely occupied by a magnesium ion. Both residues D170 and E122, which interact with the magnesium ion, play critical roles in ATP-mediated receptor activation. Mutating either residue to alanine significantly reduced the potency and affinity of α,β-methylene ATP (Table 1), indicating the importance of this metal ion site in ATP-mediated P2X1 receptor activation.

### NF449 binding site of the P2X1 receptor

NF449, a large polar molecule, is recognised for its high affinity and potency for the P2X1 receptor.^24^ The activity of NF499 was tested using a competition radioligand binding assay and calcium influx assay, demonstrating a binding affinity of 5.3 nM (Supplementary Fig. 7) and an inhibitory potency of 66 nM, respectively (Table 2). The decrease in inhibitory potency observed compared to the previously reported value of 0.05 nM at human WT-P2X1 receptors could be attributed to the utilisation of the voltage clamp technique on Xenopus oocytes as opposed to cell assays on HEK293 cells expressing the P2X1 receptor.^24^ As mentioned, NF449 adopts a distinctive orientation, requiring substantial movement of residues within the binding site, and engages in numerous interactions with the P2X1 receptor within the binding site (Fig. 5A). The P2X1 receptor engages in several salt bridge interactions with NF449, involving residues K68, K70, K136, K215, and R292 (Supplementary Table 3). Additionally, residues L72 (backbone carbonyl and amine), K140 (backbone amine), M214, C217 (backbone amine), and N290 form hydrogen bonds with NF449. The multitude of highly polar interactions, particularly around the sulfonic acids of NF449, likely underpins its high affinity for the P2X1 receptor. F188 engages in a pi-pi stacking interaction with the one benzene ring of NF449, while R139 participates in a pi-cation interaction at another benzene ring. Six additional residues, T186, V209, C217, L218, P228, and V229, were found in close proximity to NF449, suggesting their involvement in hydrophobic interactions, typically around the benzene rings of NF449.

**Figure 5.**
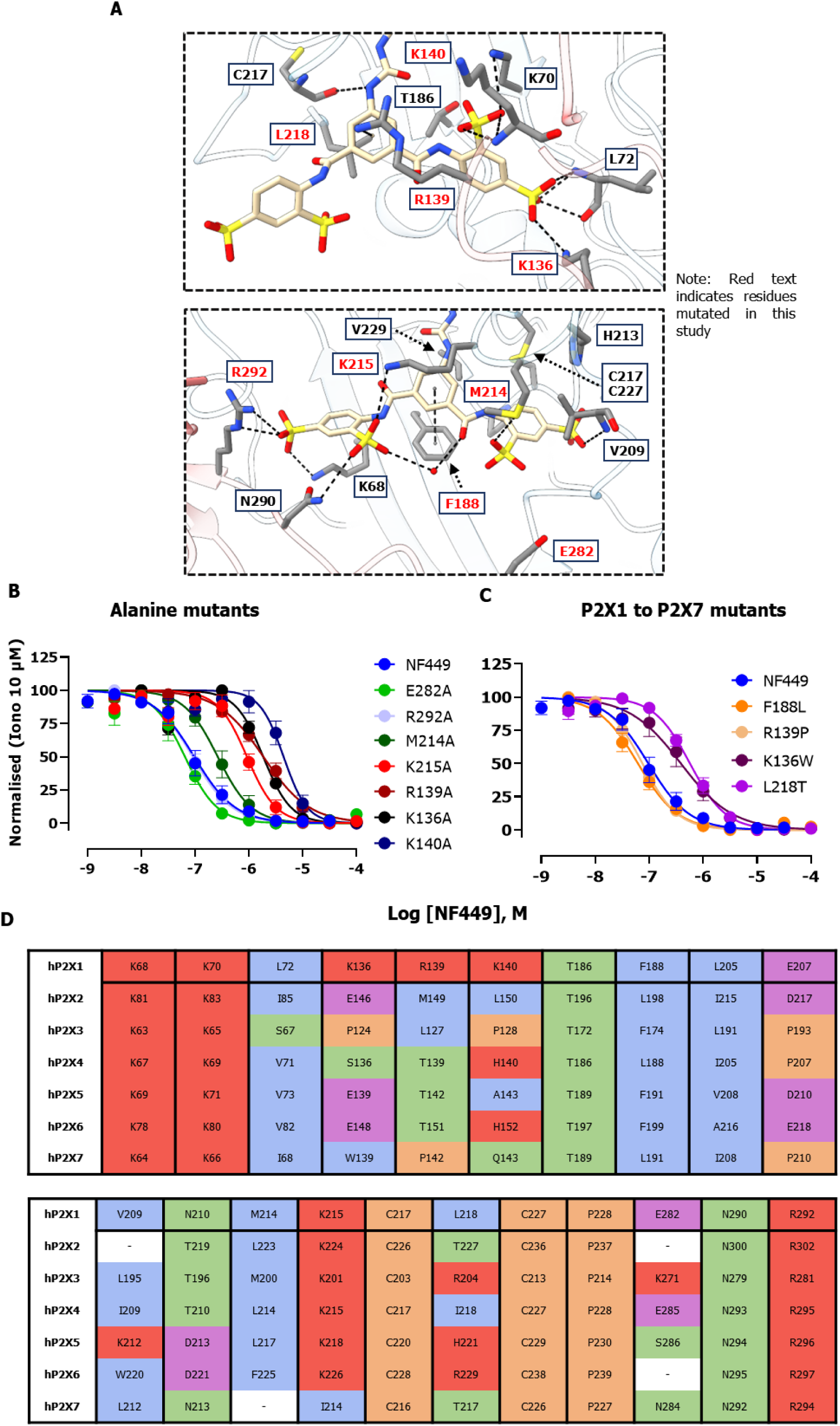
**A)** Depiction of the NF449 binding site where the key interactions are illustrated as black dotted lines, accompanied by labelling of nearby residues. **B, C)** Submaximal concentrations of α,β-methylene ATP (0.316 - 100 µM, 80-95% of max per mutant) with increasing concentrations of the antagonist NF449 on HEK293 cells expressing WT-P2X1 and single residue mutants of the P2X1 receptor. The data is normalized to 10 µM ionomycin and subsequently adjusted so that the highest value corresponds to 100 and the lowest value to 0. It is then fitted to a log(antagonist) nonlinear regression curve fit (four parameters) with the top and bottom constrained to 100 and 0, respectively. Results are presented as mean ± SEM, with n = 3 - 4 replicates. The left graph contains data for single alanine mutants, and the right graph contains data for P2X1 to P2X7 single residue mutants. **D)** Amino acid sequence alignment of key residues from the human P2X1 receptor (hP2X1:P51575), aligned with corresponding residues from other human P2X receptor subtypes (hP2X2:Q9UBL9, hP2X3:P56373, hP2X4:Q99571, hP2X5:Q93086, hP2X6:O15547, hP2X7:Q99572). Residue categorisation based on properties: Hydrophobic (A, I, L, M, F, W, V, Y) in blue, positive charge (K, R, H) in red, negative charge (E, D) in magenta, polar (N, Q, S, T) in green. Special cases (C, G, P) in orange. Gaps are represented in white.

**Table 2.** Calculated IC50 values of NF449 derived from pooled data of three to four individual experiments from HEK293 cells expressing WT-P2X1 and single residue mutants of the P2X1 receptor, alongside their respective 95% confidence intervals. Values are coloured green if within half a log unit of WT-P2X1 and coloured orange if over half a log unit of WT-P2X1 with all values over a half log unit statistically significant as calculated by a Dunnett’s multiple comparisons test.

| Receptor Type | NF449 IC50 (nM, 95% CI) |
| --- | --- |
| **P2X1 (WT)** | 66.1 (28.8 – 152) |
| **Alanine mutants** |  |
| **E282A** | 61.6 (21.5 – 177) |
| **R292A** | 91.2 (46.9 – 177) |
| **M214A** | 290.4 (73.11 – 1151) |
| **K215A** | 1011.6 (632.4 – 1622) |
| **R139A** | 2148 (922.6 – 4989) |
| **K136A** | 2317 (1439 – 3741) |
| **K140A** | 5508.1 (2624.2 – 11561) |
| **P2X1 to P2X7 mutants** |  |
| **F188L** | 56.2 (25.5 – 124) |
| **R139P** | 58 (34 – 98) |
| **K136W** | 427.6 (112.7 – 1622) |
| **L218T** | 625.2 (291.7 – 1343) |

To validate the binding mode of NF449, P2X1 receptor residues, M214, K140, E282, R292, K215, R139, and K136, were mutated to alanine. To explore the molecular basis behind the high selectivity of NF449 for the P2X1 receptor,^36^ residues, F188, K136, L218, R139 were mutated to their equivalent residues in the P2X7 receptor (Fig 5D), a receptor with over 500-fold lower potency for NF449.^24^ Initially, the potency and binding affinity of α,β-methylene ATP to each mutant were validated to ensure the mutant was functional. The mutated P2X1 receptor M214A, R139A, R139P, and R292A showed reduced potency (Table 1, Fig. 4C) and affinity (Supplementary fig. 5) for α,β-methylene ATP. The residues R292, R139, and M214, as previously discussed, interact with ATP. Furthermore, R292 plays a crucial role by forming a salt bridge with the gamma phosphate of ATP (Fig 4A). This underscores the significance of these residues in receptor activation. Conversely, P2X1 receptor mutants K215A, E282A, K136A, F188L, K136W and L218T, which do not make contacts with bound ATP, maintained similar potency (Table 1, Fig. 4C) and affinity (Supplementary fig. 5) towards α,β-methylene ATP comparable to that of WT-P2X1. To ensure similar occupancy of each mutant receptor, an 80 to 95% maximal concentration of α,β-methylene ATP was used when testing increasing concentrations of NF449 in competition binding assays. Compared to WT-P2X1, mutants M214A, K215A, R139A, K136A, and K140A exhibited a significant 4-fold, 15-fold, 32-fold, 35-fold, and 83-fold decrease in the inhibitory potency of NF449 respectively (Table 2). Residue E282, which is nearby NF449, but does not directly interact with NF449, yielded no change in the potency of NF449 when mutated to alanine (Table 2). The residues K215, R139, K136, K140, and M214 in the P2X1 receptor, crucial for NF449 activity, participate in salt bridge or hydrogen bond interactions with NF449. These interactions highlight the significance of at least four sulfonic acids and a benzene ring in NF449, underscoring their crucial role in the inhibitory activity of NF449 (Fig 4A). Furthermore, these interactions take place at various locations within NF449, indicating the accuracy of the binding mode of NF449 in the P2X1 receptor structure.

To investigate the molecular components that dictate NF449 selectivity for the P2X1 receptor four single residue mutants of the P2X1 receptor, F188L, K136W, L218T, and R139P (Fig. 4A) were created by substituting the corresponding residue from the P2X1 receptor with the sequence-matched residue from the P2X7 receptor (Fig. 5D). Among these mutants, K136W and L218T of the P2X1 receptor significantly attenuated the inhibitory activity of NF449, while F188L and R139P showed no observable effect (Fig 5C, Table 2). Residue K136 formed a salt bridge with a sulfonic acid of NF449, and L218 formed hydrophobic contacts with one of the aromatic rings of NF449. Residue F188 engages in pi-pi stacking with one of the aromatic rings of NF449. Surprisingly, mutating it to leucine (F188L), expected to disrupt this interaction, did not significantly affect NF449’s affinity suggesting that either this interaction isn’t particularly strong or that leucine may retain some hydrophobic interactions with NF449. Surprisingly, the R139P mutation did not alter the inhibitory potency of NF449 even though residue R139 appears to form a pi-cation bond with one of the aromatic rings of NF449 (Fig. 5A). However, the R139A mutant significantly reduced NF449’s inhibitory potency, indicating that the proline mutation may have retained interactions with NF449, while the alanine mutation abolishes an interaction necessary for NF449’s inhibitory activity. These results suggest that specific residues contribute to NF449’s selectivity for the P2X1 receptor over other P2X receptor subtypes.

### Comparing the orthosteric binding site at the P2X receptors

The cryo-EM structure of the ATP-bound P2X1 receptor revealed that ATP exhibits a remarkably similar binding mode to the ATP-bound P2X3, P2X4 and P2X7 receptor structures. Alignment of these structures shows that ATP from the P2X3, P2X4, and P2X7 receptors deviate by a root mean square deviations (RMSD) of 0.70, 0.93, and 1.3 Å, respectively (Fig. 6). Moreover, the interactions of bound ATP in the P2X1 receptor closely resemble those in the ATP-bound P2X3, P2X4, and P2X7 receptor structures (Fig. 6). Despite these consistent interactions across ATP-bound P2X receptor structures, it is surprising that the affinity of ATP for the P2X receptor can vary between these subtypes.^5^ The affinity of ATP for the P2X1 receptor ranks among the highest compared to other P2X receptors. Upon examining interactions of ATP at the P2X1 receptor, it became apparent that there are additional interactions, which may account for this high affinity (Fig. 6). The residues K140, D170, E122 and R139 each interact with bound ATP in the P2X1 receptor with residue K140 forming a hydrogen bond with the ribose sugar, a feature not observed in other P2X receptors. Similarly, residues D170 and E122 interact with the adjacent magnesium ion or the water network associated with the magnesium ion, while R139 interacts with the adenine ring of ATP through a water molecule. The mutagenesis data in this study highlighted the significance of D170, E122, and R139 in receptor activation (Table 1). However, it is the combination of additional interactions that likely contributes to the increased affinity of ATP for the P2X1 receptor.

**Figure 6.**
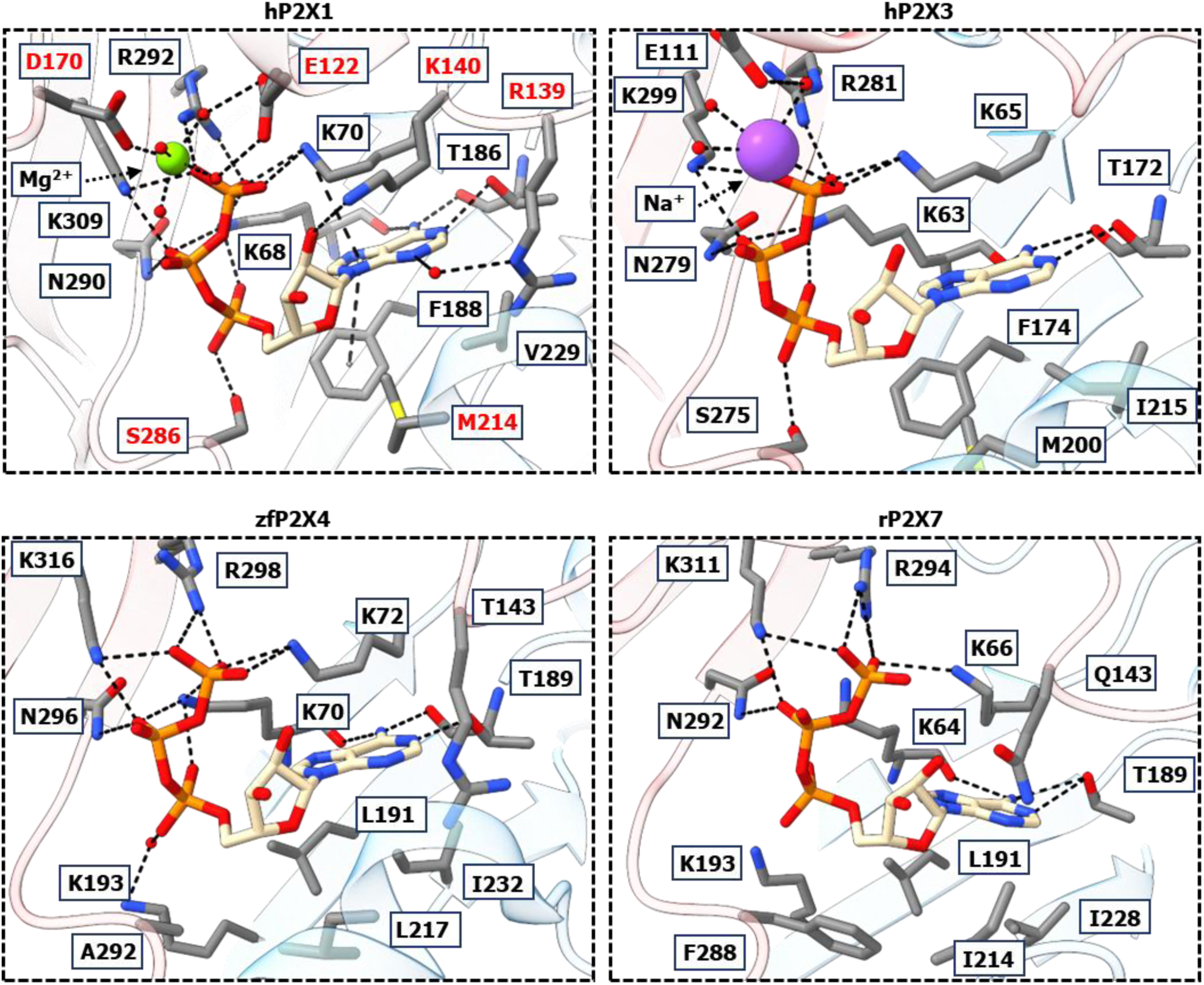
Magnified image of the ATP binding site at the P2X1, hP2X3 (5SVL), zfP2X4 (4DW1), and rP2X7 (6U9W) receptor structures where the key interactions are illustrated as black dotted lines, accompanied by labelling of nearby residues. RMSD of ATP in the P2X1 receptor compared to bound ATP in the hP2X3, zfP2X4, and rP2X7 receptors: 0.70, 0.93, and 1.3, respectively.

To explore the binding mode of NF449, P2X receptor structures bound to orthosteric antagonists were aligned with the NF449-bound P2X1 receptor structure (Fig. 7). The antagonists A-317491 from the hP2X3 (5SVR), TNP-ATP from the ckP2X7, (5XW6) and PPNDS from the pdP2X4 (8JV8) receptor structures all overlap with the ATP binding site and a portion of the NF449 binding site.^11,37, 38^ Notably, A-317491 has additional interactions at the dorsal fin of the P2X1 receptor, similar to NF449. A-317491 was designed to be a specific P2X3 receptor antagonist and does retain some weak activity for the P2X1 receptor.^39^ However, derivatives of NF449, including those featuring NF449 split in half on its symmetrical axis, exhibited large decreases in potency compared to NF449, suggesting that a large portion of NF449 is required to retain activity at the P2X1 receptor.^25^

**Figure 7.**
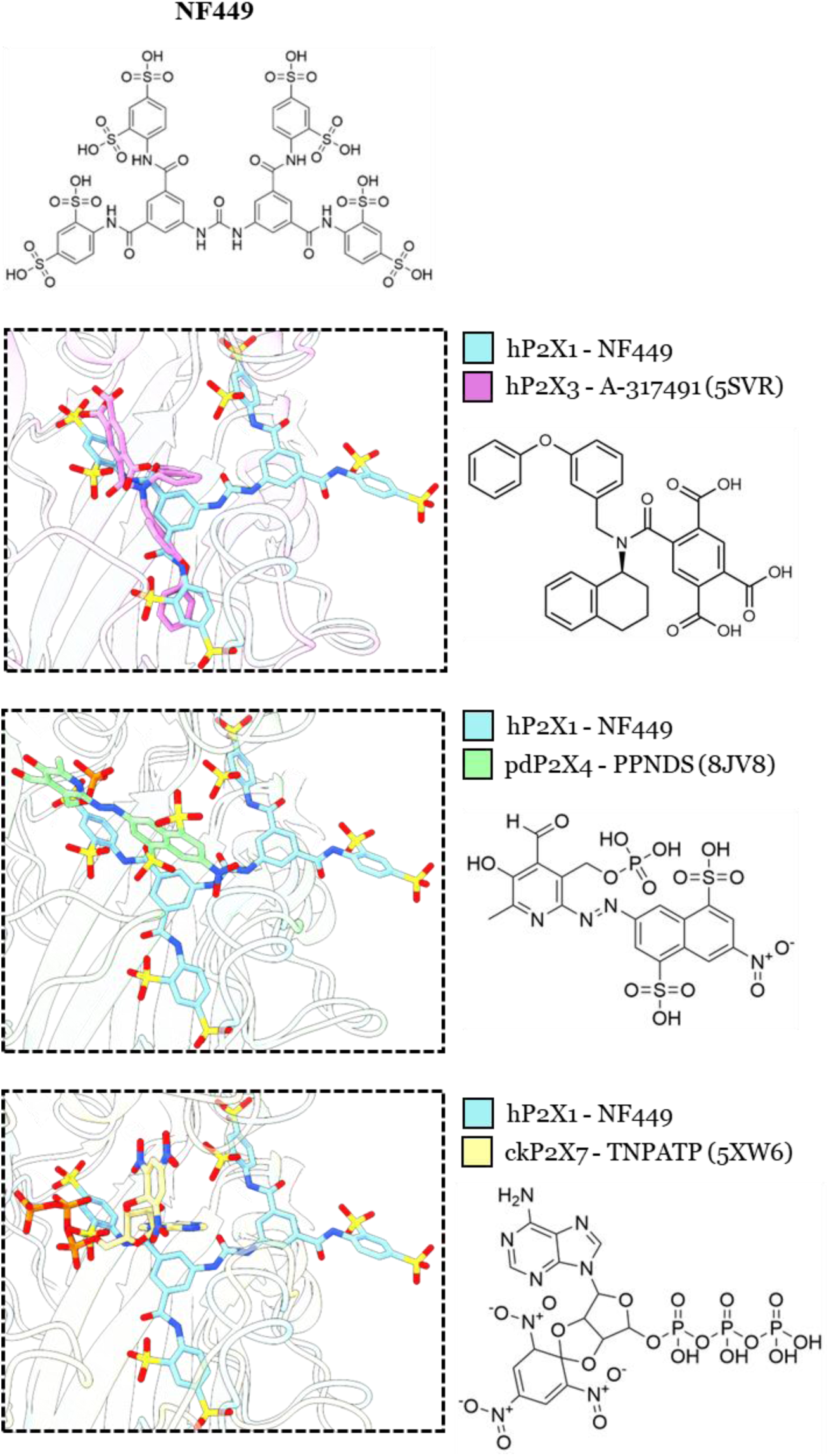
Overlay of the NF449 binding site from the P2X1 receptor structure (blue) compared to A-317491 from the hP2X3 receptor (5SVR, violet), PPNDS from the pdP2X4 receptor (8JV8, green), and TNP-ATP from the ckP2X7 receptor (5XW6, yellow). The 2D chemical structures for each of these ligands are depicted.

## Discussion

This study presents high-resolution cryo-EM structures of the P2X1 receptor in a desensitised ATP-bound and closed NF449-bound state. Features such as the binding site and gating cycle were studied in relation to other P2X receptors. Additionally, mutagenesis was used to study the interactions of ATP and NF449 and uncovered conserved and non-conserved residues that govern the binding of NF449 and ATP.

The magnesium ion adjacent to ATP was well described by metal binding parameters (Supplementary Fig. 8). However, this metal ion site is not exclusive to the P2X1 receptor, as a comparable site was identified in the crystal structures of the human P2X3 receptor and the Gulf Coast tick P2X receptor.^11,40, 41^ The functional and structural significance of this site was previously elucidated by comprehensive structural and functional studies predominantly focusing on the closely related P2X3 receptor.^38,40, 42^ It was demonstrated that the P2X3 receptor exhibits two cation binding sites within a highly acidic pocket located adjacent to the ATP binding site. It was also shown that ATP complexed with magnesium improves the affinity of ATP for the P2X3 and P2X1 receptor, while P2X2 and P2X4 receptors display notably diminished responses to Mg-ATP.^42^ In this study, mutagenesis studies and structural data demonstrate the importance of the ionic bond between D170 and the magnesium ion, as well as E122’s interaction with the water network surrounding the magnesium ion, in ATP-mediated P2X1 receptor activation (Table 1, Fig 4A). These findings corroborate previous research in this area.

The cryo-EM map of the P2X1 and NF449 bound P2X1 receptor suggested there may be another metal ion site located centrally within the extracellular domain coordinated by the D97 residue from each monomer. Mutating D97 to alanine showed this site was not essential for receptor activation or agonist binding as both calcium activation and radioligand binding assays confirmed there was no difference to WT-P2X1 (Supplementary Fig. 9). In P2X receptors, P2X4, P2X5, and P2X6 possess either aspartic acid or glutamic acid at the sequence-matched residue, implying the possible presence of this site in these receptors as well (Supplementary Fig. 9E). Indeed, Kawate et al. previously reported a site coordinated by the sequence-aligned residue E98 in the closed structure of zfP2X4, suggesting its potential involvement in ATP-mediated receptor activation. However, the lack of change observed when this residue was mutated to alanine, rendering it unable to coordinate a metal ion, implies that this site may serve a different role in the P2X1 receptor (Supplementary Fig. 9C, D).

NF449 is an interesting molecule as it is highly potent and specific to the P2X1 receptor yet, it is a poor drug candidate due to its off-target effects and high polarity.^26,36^ Mutagenesis of the NF449 binding site demonstrated that sections of bound NF449, particularly at least four sulfonic acids and a benzene are important for its inhibitory activity (Fig. 5A, Table 1). It is noteworthy that R292, a residue that forms a salt bridge interaction with the P2X1 receptor, did not exhibit any change in the inhibitory activity of NF449 when mutated to alanine (Table 2). Furthermore, some sections of NF449 do not form any interactions with the P2X1 receptor (Fig 5A), indicating this molecule could be further optimised. This underscores the need for a more drastic modification of the chemical structure of NF449 to improve drug properties while retaining activity. It remains to be seen if the unique binding mode of NF449 could be optimised to produce a more drug-like molecule.

The conserved nature of the orthosteric pocket across P2X receptors prompts exploration of allosteric sites. Allosteric sites offer an alternative druggable site, which may be more receptor-specific. Another intriguing aspect of the NF449 binding site is the region outside of the orthosteric pocket occupied by bound NF449 which could be targeted by smaller ligands as an allosteric site (Fig. 5A). While no conclusive allosteric sites have been identified for the P2X1 receptor, suggestions have been made that recently discovered P2X1 receptor antagonists, ATA and PSB-2001, may bind to an allosteric site at the P2X1 receptor.^20,22^ One allosteric site has been identified for the P2X3 receptor and another allosteric site identified in the P2X4 and P2X7 receptors.^6,7, 43^ Comparatively, these allosteric pockets are more occluded when compared to the equivalent region in the P2X1 receptor (Supplementary Fig. 10). Furthermore, the ligands bound at these sites AF-219, BX430 and JNJ47965576 and related ligands have minimal activity at the P2X1 receptor.^44^ This indicates that new chemical scaffolds may be necessary to target these two allosteric sites effectively on the P2X1 receptor. Whether these sites will be conducive to small molecule drug discovery at the P2X1 receptor remains to be seen. Nevertheless, identifying new binding sites at the P2X1 receptor that can modulate receptor function can only be beneficial in opening new opportunities for drug discovery.

## Conclusion

The high-resolution structures of the desensitised ATP-bound and closed NF449-bound P2X1 receptors unveil novel structural features and essential interactions, shedding light on the receptor architecture and its distinctive binding mechanism with the potent antagonist NF449. These structural revelations open exciting opportunities for innovative structure-based discovery efforts in the realm of P2X1 receptor modulation. As the development of new therapeutics targeting the P2X1 receptor is still in its early stages, these structures are poised to guide the design of novel tool compounds, which may play a pivotal role in advancing our understanding and harnessing the therapeutic potential of the P2X1 receptor.

## Methods

### Expression and purification of the P2X1 receptor

The full-length human P2X1 gene with a C-terminal tag containing a 3C-cleavage site followed by muGFP and an octa-histidine sequence (Supplementary Fig. 1A), were cloned into the pFastBac vector. Utilising the Bac-to-Bac system, the P2X1 construct was transformed into a bacmid to make baculovirus and transfected into *S. frugiperda* cells using FuGENE HD transfection reagent. The P2X1 receptor was then expressed in *S. frugiperda* cells using baculovirus, with cell harvest conducted approximately 60 hours post-transfection, followed by freezing at −80⁰C.

Purification protocols were initially derived from existing literature reports on P2X receptor purifications and subsequently optimized specifically for the P2X1 receptor. Cell pellets were thawed and lysed using a buffer consisting of 50 mM Tris (pH 8), 1 mM EDTA, 1 mM MgCl_2_, 2.5 µL/L benzonase, and a protease inhibitor cocktail containing, 0.2 mM phenylmethylsulfonyl fluoride, 5 µg/ml leupeptin, 5 µg/ml soybean trypsin inhibitor, and 0.2 mg/ml benzamidine. The resulting cell membranes were then subjected to centrifugation and subsequently resuspended in a solubilization buffer comprising 50 mM Tris (pH 8), 15% glycerol, 750 mM NaCl, 1 mM MgCl, 5 mM imidazole, 2.5 µL/L benzonase, 0.5% Lauryl Maltose Neopentyl Glycol (LMNG), 0.03% Cholesteryl Hemisuccinate Tris Salt (CHS), and the protease inhibitor cocktail. The cell membranes were homogenized using a dounce and stirred for 1.5 hours. Following this, the sample underwent high-speed centrifugation and was then batch bound to pre-equilibrated Talon resin for 2 hours. The sample was subsequently washed with a buffer containing 50 mM Tris (pH 8), 5% glycerol, 750 mM NaCl, 20 mM imidazole, 0.01% LMNG, and 0.0006% CHS. Upon loading onto a glass column, it was washed with 6 column volumes, and elution was performed using elution buffer comprising 50 mM Tris (pH 8), 5% glycerol, 100 mM NaCl, 200 mM imidazole, 0.01% LMNG, and 0.0006% CHS, collecting 5 column volumes. The sample underwent buffer exchange through a Amicon Ultra-15 100 KDa molecular mass cut-off centrifugal filter unit (Millipore, USA), followed by cleavage with 3C protease at a 1:1 w/w ratio overnight at 4⁰C. The cleaved sample was then passed through fresh Talon resin, and the flowthrough was collected. Subsequently, size exclusion chromatography (SEC) was employed for purification, utilizing SEC buffer consisting of 50 mM Tris (pH 8), 100 mM NaCl, 0.01% LMNG, and 0.0006% CHS. The sample was loaded onto a pre-equilibrated Superdex 200 Increase 10/300 GL column (Cytiva, USA), and fractions containing the trimeric P2X1 receptor were pooled, concentrated, flash-frozen in liquid nitrogen, and stored at −80⁰C. This purification process yields the ATP-bound desensitised P2X1 receptor as determined by cryo-EM studies.

To obtain the NF449-bound P2X1 receptor, the aforementioned protocol is modified with the inclusion of 100 µM NF449, 5 µL of apyrase, and 500 µM CaCl2 during the overnight 3C cleavage at 4⁰C. Additionally, 1-2 µL of apyrase is incorporated into both the elution and Ni wash buffers.

To verify sample purity and homogeneity, each sample was loaded onto TGX precast (4% - 15%) polyacrylamide gels (Bio-rad, USA) alongside Precision Plus Protein Dual Color Standards. Following electrophoresis, gels were stained with Coomassie blue and imaged with a camera. Additionally, samples were validated through negative stain electron microscopy with uranyl formate. Stained grids were visualized using a Talos L120C 120kV microscope (Thermo Fisher Scientific, USA).

### Single particle cryogenic transmission electron microscopy

To determine the structure of the ATP-bound P2X1 receptor, purified ATP-bound P2X1 receptor (19 mg/ml) was supplemented with 1 mM ATP and 1 mM MgCl2 overnight before vitrification. Additionally, 0.4 mM or 1.2 mM of fluorinated Fos-Choline-8 (fluor-FC8) was added just one minute prior to vitrification as conducted in previous studies.^45^ To determine the structure of the NF449-bound P2X1 receptor, purified NF449-bound P2X1 receptor (15.5 mg/ml) was supplemented with 100 µM NF449 overnight prior to vitrification. Additionally, 0.3 mM of fluorinated Fos-Choline-8 (fluor-FC8) was added one minute prior to vitrification. All samples were pipetted at a volume of 3 µL onto UltrAufoil R1.2/1.3 300 mesh holey grids (Quantifoil, Germany) which had been glow discharged in air at 15 mA for 180 s using a Pelco EasyGlow. Grids were plunge frozen in liquid ethane using a Vitrobot Mark III (Thermo Fisher Scientific, USA) operated at 4⁰C and 100% humidity with 12 blot force and 2 seconds blot time. The cryo-EM data were acquired at the Ramaciotti Centre for Cryo-Electron Microscopy, on a G1 300 kV Titan Krios microscope (Thermo Fisher Scientific, USA) fitted with S-FEG, a BioQuantum energy filter and K3 detector (Gatan, Pleasanton, California, USA). The Krios microscope was operated at an accelerating voltage of 300 kV, utilizing a 50 μm C2 aperture, and a 100 µm objective aperture, along with zero-loss filtering with a slit width of 10 eV. The microscope was set to an indicated magnification of 105,000 times in nanoprobe EFTEM mode. Data collection was assisted using aberration-free image shift with Thermo Fisher Scientific EPU software. Additional microscope details are listed in supplementary table 1.

### Cryo-EM image processing and model building

The workflow for determining the ATP-bound and NF449-bound P2X1 receptor structures is detailed in Supplementary Figure 2. Movies were collected at 0.82 Å per pixel and processed through motion correction and CTF correction using CryoSPARC v3-4.^46^ Particle picking was performed using template picking, followed by multiple rounds of 2D and 3D classification within CryoSPARC v3-4. Bayesian polishing was applied to the ATP-bound P2X1 receptor in RELION v3.1.^47^ A CryoSPARC 3D variability analysis was conducted on a larger set of NF449-bound P2X1 receptor particles in clustered mode (2 clusters). This analysis was used to eliminate a subset of particles representing a different state (desensitised) from the expected state (closed). The final homogenous refinement for the ATP-bound receptor utilised 481,309 combined particles from two grids, resulting in a 1.96 Å map (FSC = 0.143). The NF449-bound receptor underwent non-uniform refinement, yielding a 2.61 Å map (FSC = 0.143) from 41,932 particles. Both maps were refined with C3 symmetry applied. Initial models were obtained from the AlphaFold Protein Structure Database^48,49^ and rigid body fitted into the density using UCSF ChimeraX.^50^ Multiple rounds of manual model building and real-space refinement were conducted in Coot^51^ and PHENIX^52^, respectively. Residues in the intracellular region of the ATP-bound P2X1 receptor (M1-G30 and L357-S399) and the NF449-bound P2X1 receptor (M1-G30 and D350-S399) were not modelled due to poor density in the cryo-EM map. Model quality was assessed using MolProbity^53^ before PDB deposition. Structure figures were generated using ChimeraX v1.6. Ligand interactions and metal ion analysis were performed using ChimeraX v1.6 and CheckMyMetal: metal binding site validation server^31^, respectively. Pore radius calculation was conducted in Mole v2.5^54^ and visualized in ChimeraX v1.6.

### P2X1 receptor stable transfection in mammalian cells

WT-P2X1 or single-residue mutants of the P2X1 receptor were cloned into pcDNA5/FRT/TO. These constructs were stably transfected into HEK293 Flp-In T-Rex cells. Transfection involved a FuGENE mixture composed of 250 µL Opti-MEM Reduced Serum Medium, 0.7 µg P2X1 construct, 6.3 µg pOG44 (in a 10:1 ratio to P2X1 construct), and 28 µL FuGENE (in a 4:1 ratio to total DNA), incubated for 30 minutes before addition to cells. After 48 hours, selection with 175 µg/mL Hygromycin B was initiated until individual colonies formed. These colonies were reseeded, expanded under selection conditions, and preserved by freezing. To induce P2X1 receptor expression tetracycline was added at a concentration of 150 ng/mL two days prior to experimental testing.

### Radioligand binding assays

HEK293 cells expressing either WT-P2X1 or single residue mutants of the P2X1 receptor were seeded in 96-well isoplates at a density of 15,000 cells per well. For the experiments, a buffer solution with the following composition: 150 mM NaCl, 2.6 mM KCl, 1.18 mM MgCl2.6H2O, 10 mM D-glucose, 10 mM HEPES, 2.2 mM CaCl2.H2O at pH 7.4 was prepared, and the cells were washed twice. Saturation assays involved the addition of increasing concentrations of [^3^H]-α,β-methylene ATP (Moravek Inc., USA). Competition assays utilized a sub-saturation concentration of [^3^H]-α,β-meATP (50 nM), along with increasing concentrations of P2X1 receptor ligands. ATP was employed to assess non-specific binding of [^3^H]-α,β-methylene ATP in these experiments. Experiments were performed in triplicate, and assay plates were incubated at room temperature for at least two hours. Optiphase SuperMix scintillant was added to each well and plates were read using a MicroBeta counter (PerkinElmer, USA). [^3^H]-α,β-methylene ATP concentrations were checked on Tri Carb scintillation beta counters (PerkinElmer, USA). Data analysis was conducted using GraphPad Prism v9. Saturation radioligand experimental data were fitted to a one-site specific binding curve, with results reported in counts per minute (CPM). Radioligand competition experimental data were fitted to a one-site Ki curve, incorporating the concentration and affinity of [^3^H]-α,β-methylene ATP and results were reported in specific binding.

### Intracellular calcium influx (activity) assay

HEK293 cells expressing either WT-P2X1 or single residue mutants of the P2X1 receptor were seeded into 96-well clear plates at a density of 15,000 cells per well. A buffer solution was prepared, consisting of 150 mM NaCl, 2.6 mM KCl, 1.18 mM MgCl2.6H2O, 10 mM D-glucose, 10 mM HEPES, 2.2 mM CaCl2.H2O, 0.5% w/v BSA, and 2 mM probenecid acid at pH 7.4. The cells were washed twice with this buffer. Subsequently, 1 mM Fluo-8 was added to each well under reduced light conditions. The cells were then treated with ligands as required and incubated for an additional 30 minutes. For the assay, drug plates were prepared, including the non-specific calcium ionophore ionomycin, control buffer, and α,β-methylene ATP. Data were recorded using a FlexStation 3 (Molecular Devices, USA) with excitation at 485 nm and emission at 525 nm, with auto cutoff at 515 nm. Initial baseline readings were taken over a 20-second period, followed by stimulation, and measurements were recorded for an additional two minutes at regular intervals. For each condition, the maximum subtracted from the minimum data was exported and analyzed using GraphPad Prism v9. Data were normalized to 10 µM ionomycin and adjusted so that the highest value corresponded to 100 and the lowest value to 0. Agonist curves were fitted to a log(agonist) nonlinear regression curve using a four-parameter model, with the top and bottom constrained to 100 and 0, respectively. Antagonist curves were fitted to a log(antagonist) nonlinear regression curve using a four-parameter model, with the top and bottom constrained to 100 and 0, respectively. EC50 and IC50 values were extracted from WT-P2X1 or single residue mutants of the P2X1 receptor and plotted on column graphs. These data were analyzed using a one-way ANOVA Dunnett multiple comparisons test to determine if there was a significant difference (<0.05) compared to WT-P2X1.

### Kinase-Glo luminescent ATP assay

To measure ATP bound to detergent purified P2X1 receptor a Kinase-Glo® Luminescent Kinase Assay (ProMega, USA) was used. Detergent purified P2X1 receptor was incubated with apyrase to remove ATP in solution and then heated at 70⁰C for 20 minutes to release bound ATP and degrade apyrase. Samples were plated into solid white 384 well plates with a 1:1 ratio to Kinase-Glo reagent. Luminescence is recorded on a lumistar (BMG LABTECH, Germany) and graphed in GraphPad Prism v9.

## Acknowledgments

This work was funded by the National Health and Medical Research Council of Australia Investigator Grant (1196951): D.M.T). The cryo-EM imaging and sample vitrification were performed at the Monash University Ramaciotti Centre for cryo-electron microscopy. The cryo-EM data processing and data storage were done using the Monash University MASSIVE high-performance computing facility and supercomputing resources.

## Author Contributions

D.M.T., S.V., and J.I.M. designed the overall research. F.M.B. designed, expressed, and purified protein samples. F.M.B performed negative-stain EM. F.M.B., J.I.M., A.G., H.V. performed sample vitrification and cryo-EM imaging. F.M.B. processed the EM data and generated and analysed atomic models. F.M.B. generated DNA constructs and performed and analysed pharmacology experiments. D.M.T., S.V., and J.I.M. provided supervision. F.M.B. wrote the manuscript with contributions and input from all authors.

