## Supplemental Data for "Structural insights into the human P2X1 receptor and ligand interactions"

### Supplementary Figures

**Supplementary Figure 1. A)** The construct utilised for expressing the P2X1 receptor containing the full-length human P2X1 receptor along with a tag featuring a 3C cleavage site, muGFP, and an 8x His-tag, which were subsequently removed during the purification process. Purification results of the **B)** ATP-bound P2X1 receptor and **C)** NF449-bound P2X1 receptor. A size exclusion chromatography (SEC) trace was obtained using an FPLC equipped with a Superdex 200 Increase 10/300 column, with the red box highlighting the volume of the collected sample, alongside the corresponding SDS-page Coomassie-stained gel of the purified P2X1 receptor run with the Precision Plus Protein Dual Colour Standards ladder. Luminescence values from an ATP-based luciferase experiment of purified NF449-bound P2X1 receptor and ATP-bound P2X1 receptor show minimal levels of ATP in the purified NF449-bound P2X1 receptor sample.

**Supplementary Figure 2.** Imaging and processing workflow from cryosparc v3 and v4 and relion v3.1.2 for the ATP-bound P2X1 receptor and NF449-bound P2X1 receptor.

**Supplementary Figure 3.** The C3 symmetry cryo-EM maps of the **A)** desensitized ATP-bound P2X1 receptor (contour 0.29) and **B)** closed NF449-bound P2X1 receptor (contour 0.1) are presented, with local resolution indicated. Each receptor is colour-coded: dark blue represents 2 Å resolution, white indicates 3 Å resolution, and red signifies 4 Å resolution. Additionally, gold-standard Fourier shell correlation curves and conical Fourier shell correlation summary plots are displayed for the 3D reconstruction. A cFAR below 0.5 indicates the preferred orientation in the particle set. Regions of the P2X1 receptor model fit into the cryo-EM map of the P2X1 receptor with a focus on the beta sheets of the lower body, head domain and fluke (transmembrane domains).

**Supplementary Figure 4.** The C1 symmetry cryo-EM maps of the **A)** desensitized ATP-bound P2X1 receptor (contour 0.3) and **B)** closed NF449-bound P2X1 receptor (contour 0.1) are presented, with local resolution indicated. Each receptor is colour-coded: dark blue represents 2Å resolution, white indicates 3Å, and red signifies 4Å. Additionally, gold-standard Fourier shell correlation curves are displayed for the 3D reconstruction.

**Supplementary Figure 5. A)** Radioligand saturation binding with increasing concentrations of $[^3H]$ α,β-methylene ATP on HEK293 cells expressing WT-P2X1 receptor or single residue mutants of the P2X1 receptor. Data is plotted as specific binding and fit to a nonlinear regression one site – fit Ki (mean ± SEM, n = 2 - 5). **B)** Graph of grouped affinity (Ki) values of tritiated α,β-methylene ATP on HEK293 cells expressing WT-P2X1 receptor or single residue mutants of the P2X1 receptor from individual experiments (mean ± SEM, n = 2 - 5). Values that exceeded 300 nM were classified as not defined.

**Supplementary Figure 6. A)** Graph of grouped potency (EC50) values of α,β-methylene ATP on HEK293 cells expressing WT-P2X1 receptor or single residue mutants of the P2X1 receptor (mean ± SEM, n = 4). **B)** Graph of grouped affinity (Ki) values of $[^3H]$ α,β-methylene ATP on HEK293 cells expressing WT-P2X1 receptor or single residue mutants of the P2X1 receptor (mean ± SEM, n = 3 - 4). Comparisons to WT-P2X1 for each single residue P2X1 receptor mutant analysed with a one-way ANOVA Dunnett's multiple comparisons test to determine if there is a significant difference (<0.05) to WT-P2X1. Stars in the table indicate significance levels as follows: * indicates P ≤ 0.05, ** indicates P ≤ 0.01, *** indicates P ≤ 0.001, and **** indicates P ≤ 0.0001.

**Supplementary Figure 7.** Radioligand competition data of the submaximal concentration of [$^3$H] α,β-methylene ATP (50 nM) with increasing concentrations of NF449 on HEK293 cells expressing WT-P2X1 receptor. Data is plotted as specific binding and fit to a nonlinear regression one site – fit Ki (mean ± SEM, n = 3).

**Supplementary Figure 8.** Validation metrics from CheckMyMetal (CMM): Metal Binding Site Validation Server for the magnesium ion positioned adjacent to ATP, with each box colour-coded as follows: green for acceptable, yellow for borderline, and red for poor. The ATP-Mg site is modeled into the cryo-EM map density of the P2X1 receptor.

**Supplementary Figure 9. A)** Cryo-EM map and model of the ATP-bound P2X1 receptor focused on a possible central metal ion site within the extracellular domain at the threefold axis. **B)** Calculated IC50 and Ki values from WT-P2X1 and P2X1 D97A receptor-expressing HEK293 cells. **C)** Increasing concentrations of the agonist α,β-methylene ATP were applied to HEK293 cells expressing either WT-P2X1 or P2X1 D97A receptor. The data was normalized to 10 μM ionomycin, with the highest value set to 100 and the lowest value to 0. It was then fitted to a log(agonist) nonlinear regression curve using four parameters, with the top and bottom constrained to 100 and 0, respectively (mean ± SEM, n = 4). **D)** Radioligand saturation binding was conducted with increasing concentrations of tritiated α,β-methylene ATP on HEK293 cells expressing either WT-P2X1 receptor or P2X1 D97A receptor. The data was plotted as specific binding and fitted to a nonlinear regression one-site model to determine Ki (mean ± SEM, n = 2 - 5). **E)** Amino acid sequence alignment of D97 from the human P2X1 receptor (hP2X1:P51575), aligned with the corresponding residue from other human P2X receptor subtypes (hP2X2:Q9UBL9, hP2X3:P56373, hP2X4:Q99571, hP2X5:Q93086, hP2X6:O15547, hP2X7:Q99572). Residue categorisation based on properties: Hydrophobic (A, I, L, M, F, W, V, Y) in blue, positive charge (K, R, H) in red, negative charge (E, D) in magenta, polar (N, Q, S, T) in green. Special cases (C, G, P) in orange. Gaps are represented in white.

**Supplementary Figure 10.** The P2X1 receptor structure was aligned with the allosteric sites of the P2X3 (5YVE, yellow), P2X4 (8JV5, green), and P2X7 (5U1X, violet) receptors. The allosteric ligands bound in each of these receptors are represented in grey.

**Supplementary Table 1.** Summary of data collection, cryo-EM map parameters, and model validation for the ATP-bound P2X1 receptor (PDB: 9B73) and NF449-bound P2X1 receptor (PDB: 9B95).

**Supplementary Table 2.** Interactions of Mg-ATP with the P2X1 receptor from the cryo-EM structure, including hydrogen bonds (2.7 - 3.3 angstroms), salt bridges (2.7 – 4.0 angstroms), hydrophobic interactions (3.3 - 4.0 angstroms), Pi-cation and Pi-Pi interactions (3.5 - 6 angstroms), and ionic interactions with Mg2+ (1.8 - 2.5 angstroms). Interactions outside these ranges are indicated with an asterisk.

**Supplementary Table 3.** Interactions of NF449 with the P2X1 receptor from the cryo-EM structure, including: hydrogen bonds (2.7 - 3.3 angstroms), salt bridges (2.7 – 4.0 angstroms), hydrophobic interactions (3.3 - 4.0 angstroms), Pi-cation and Pi-Pi interactions (3.5 - 6 angstroms), and ionic interactions with Mg2+ (1.8 - 2.5 angstroms). Interactions outside these ranges are indicated with an asterisk.

**Supplementary Figure 1**

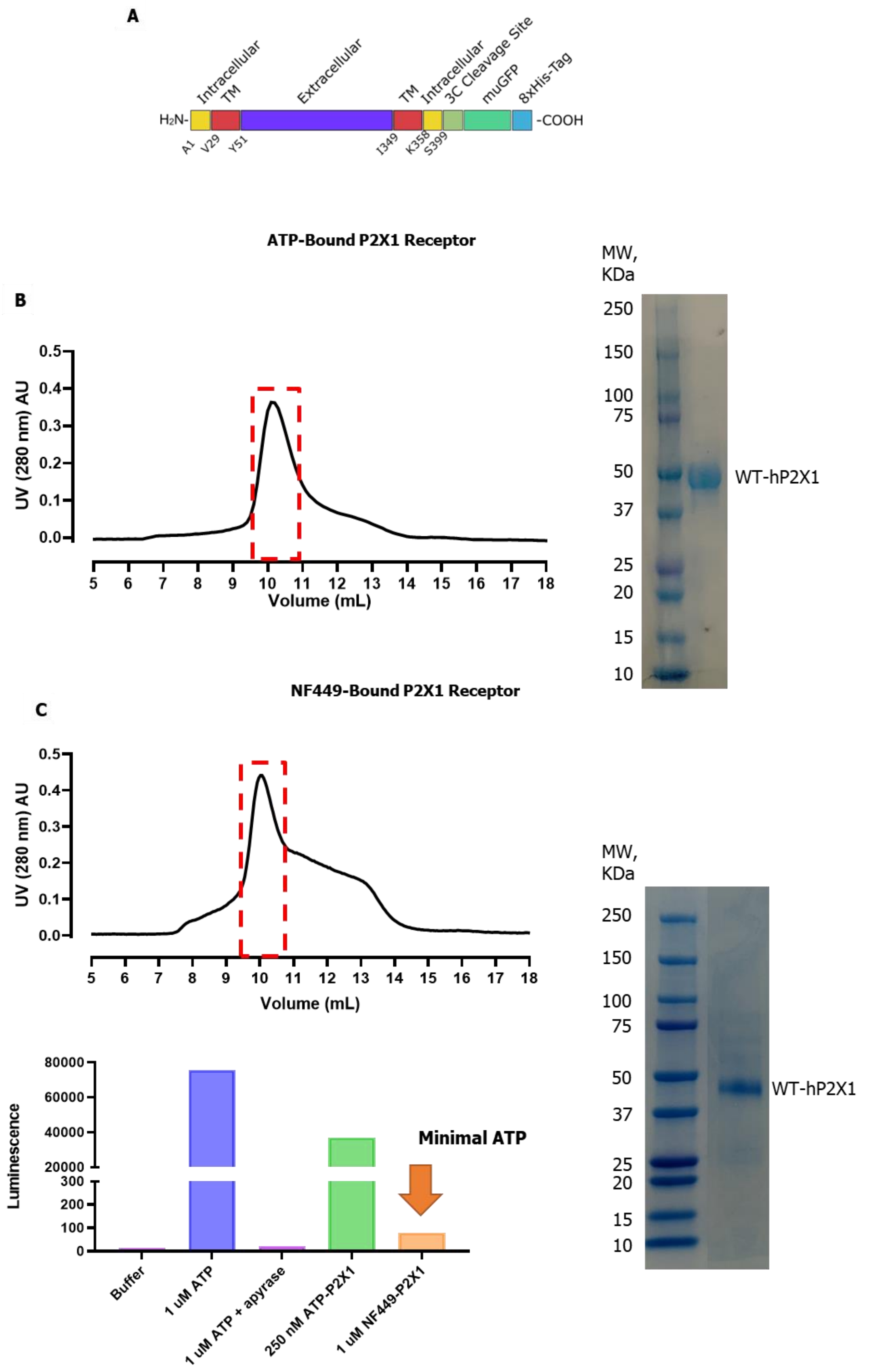

**Supplementary Figure 2**

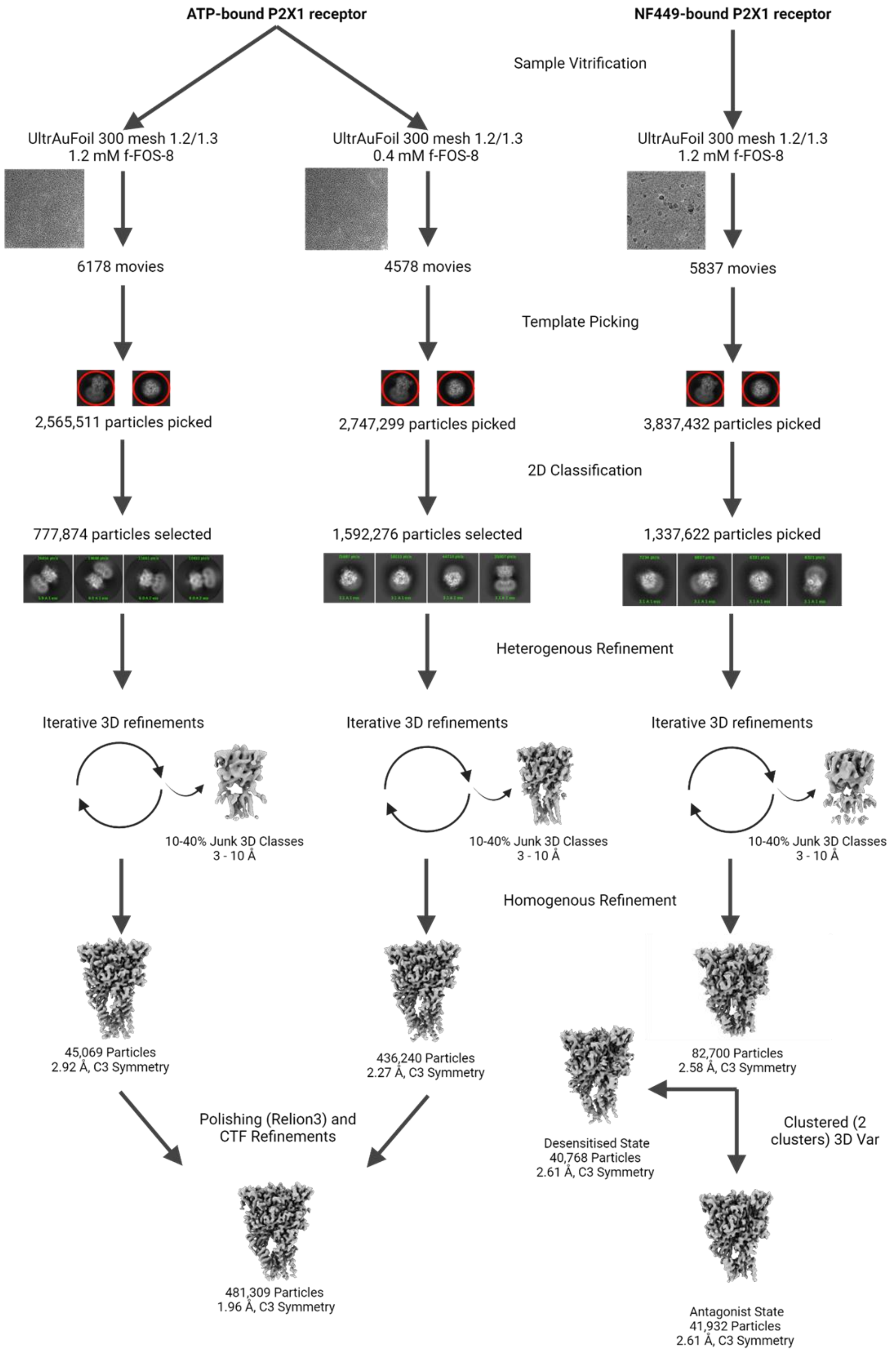

**Supplementary Figure 3**

**A**

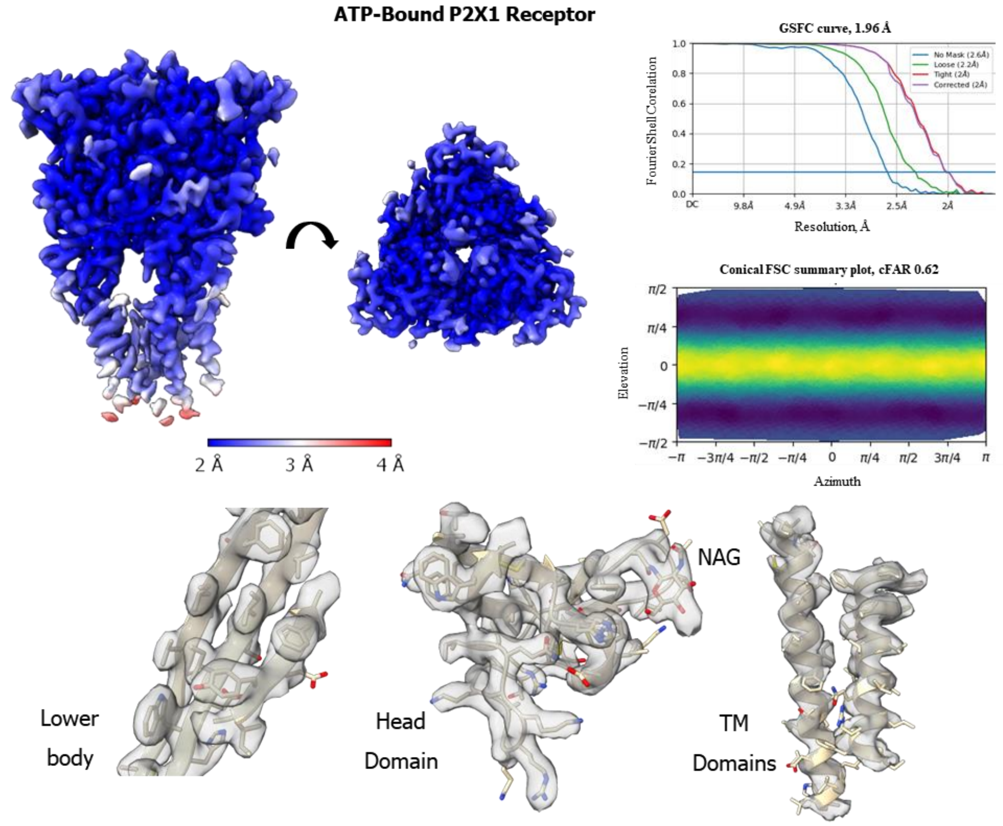


**B**

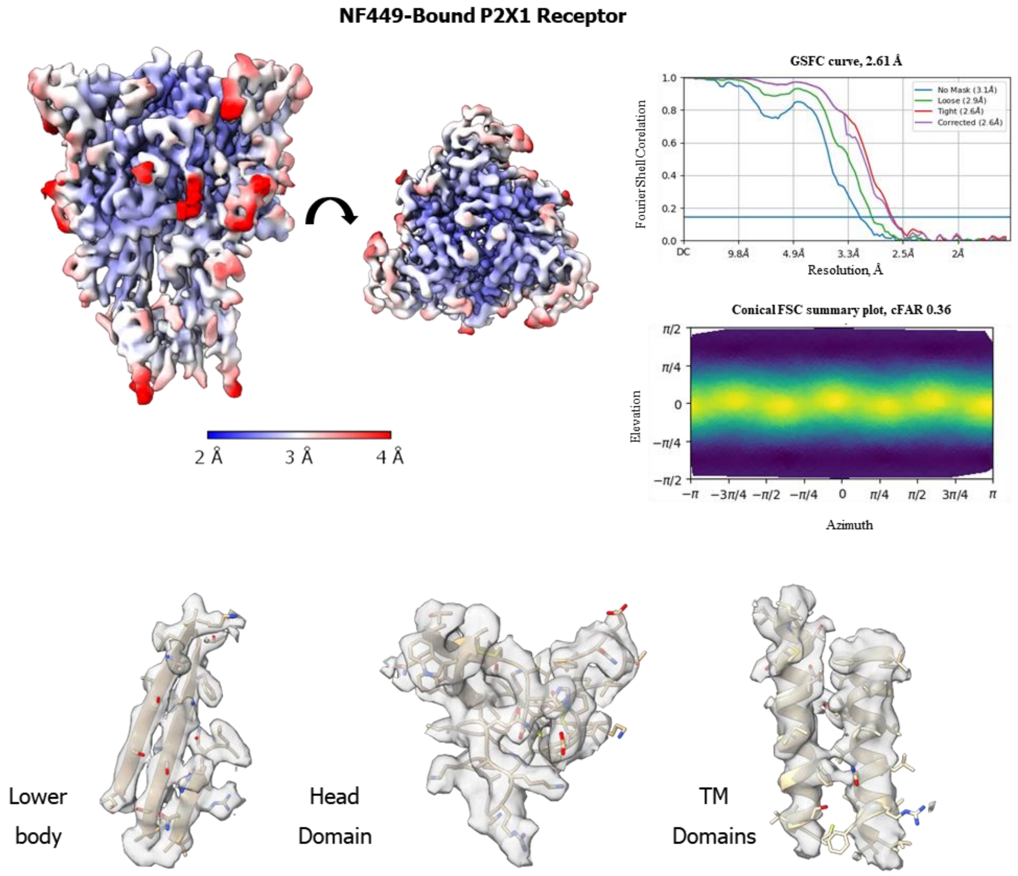

### Supplementary Figure 4

**A**

**ATP-Bound P2X1 Receptor (C1 symmetry)**

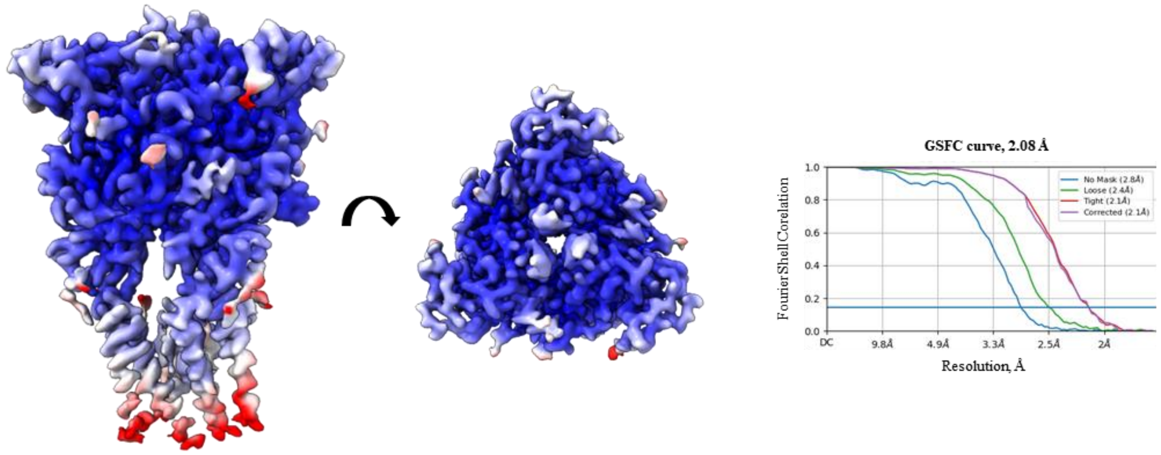


**B**

**NF449-Bound P2X1 Receptor (C1 symmetry)**

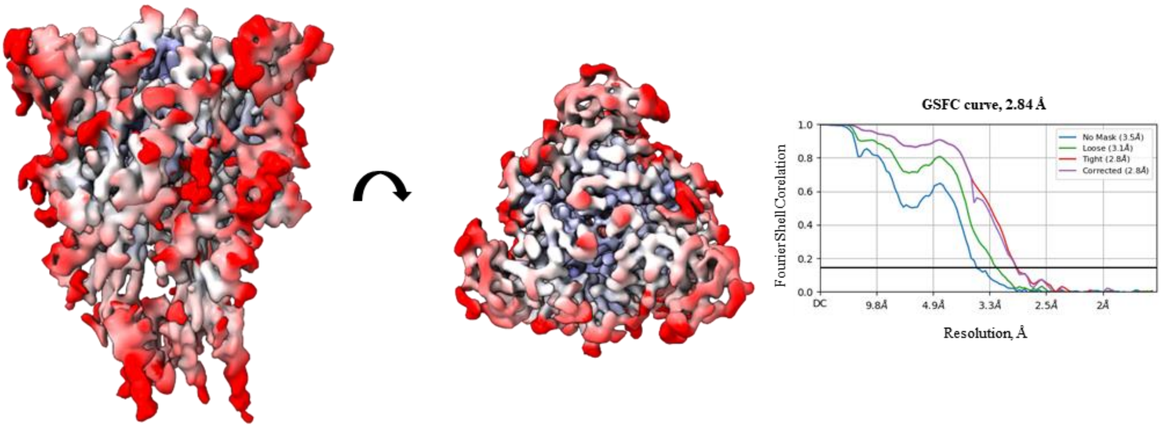

**Supplementary Figure 5**

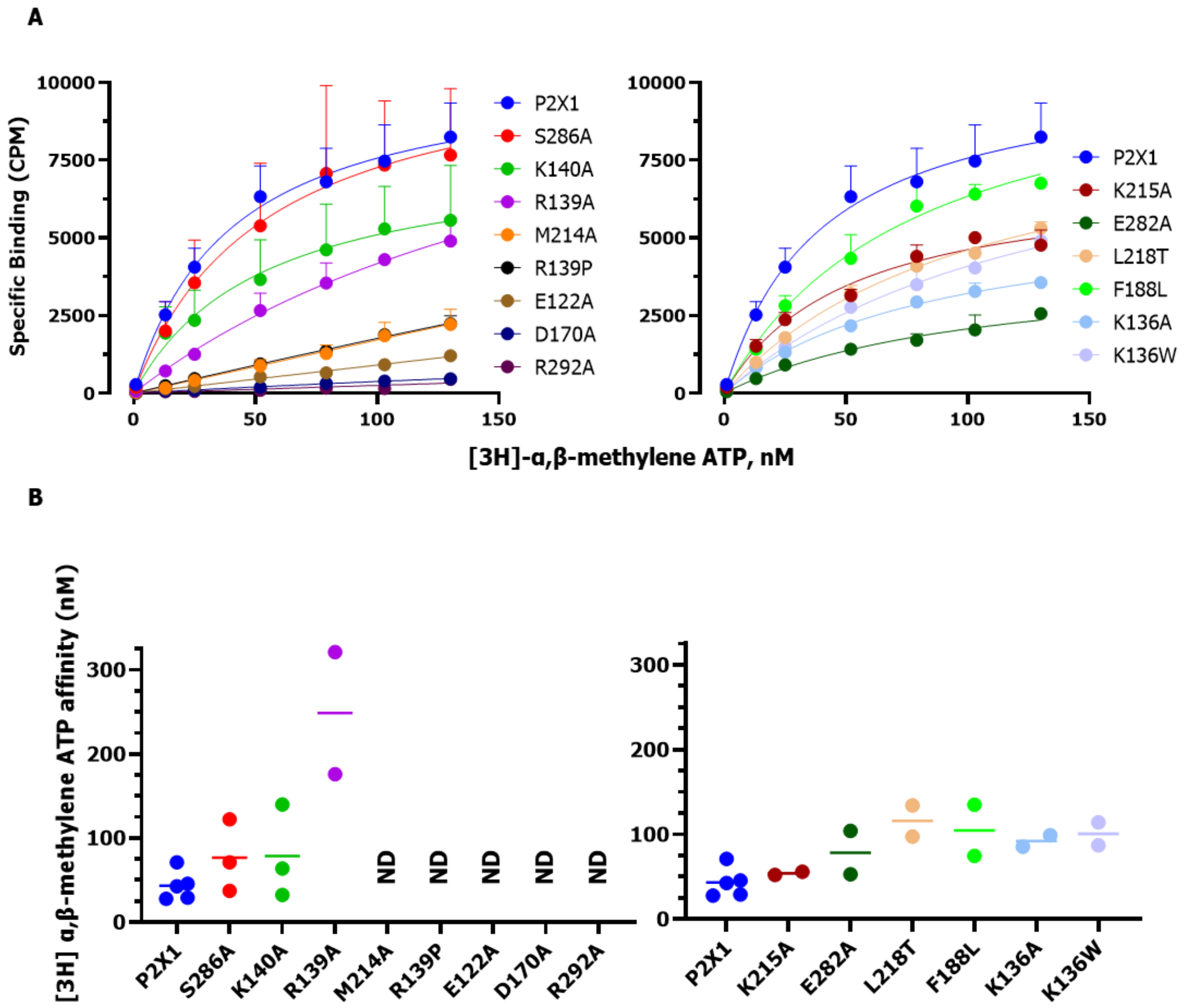

**Supplementary Figure 6**

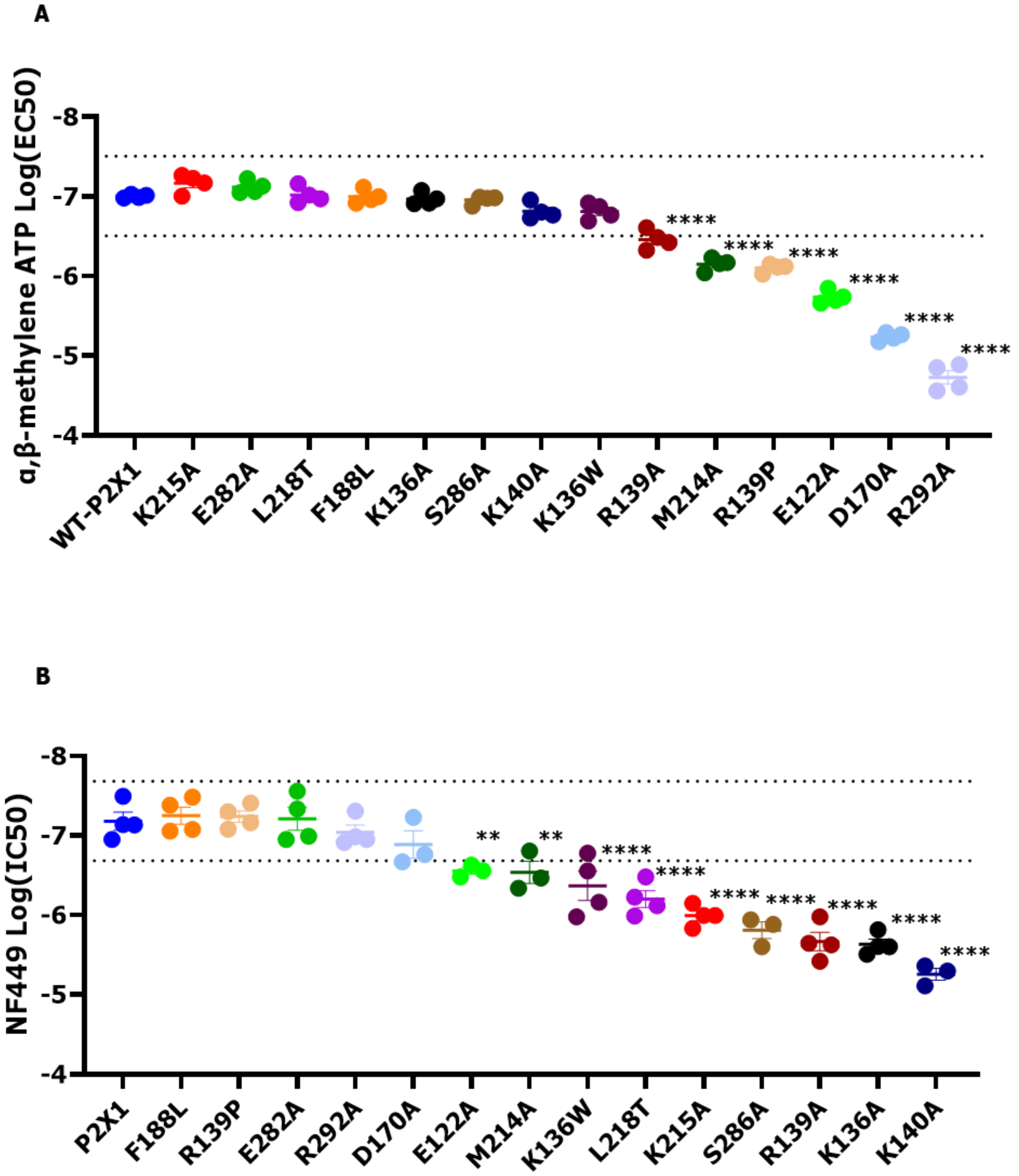

**Supplementary Figure 7**

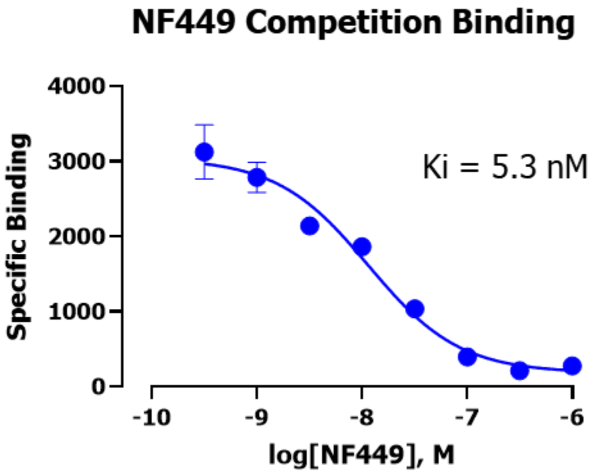

**Supplementary Figure 8**

**Mg-ATP**

| | Occupancy | B factor | Atomic Contacts | Valence | nVESCUM | Geometry | gRMSD | Vacancy |
|---|---|---|---|---|---|---|---|---|
| **Mg-ATP** | 1 | 89.9 (84.1) | $O_6$ | 2.7 | 0.16 | Octahedral | 16.1° | 0 |

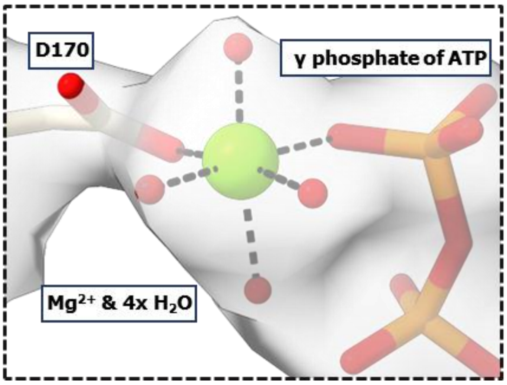

**Supplementary Figure 9**

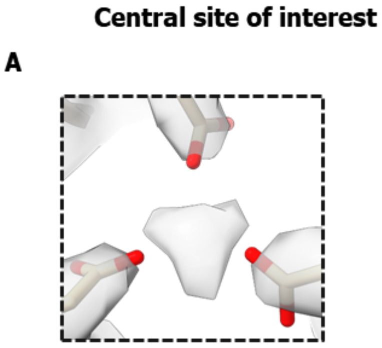


B

| Mutants | EC50 | Ki |
|---|---|---|
| P2X1 (WT) | 111.4 nM | 43.3 |
| D97A | 161.1 nM | 47.8 |

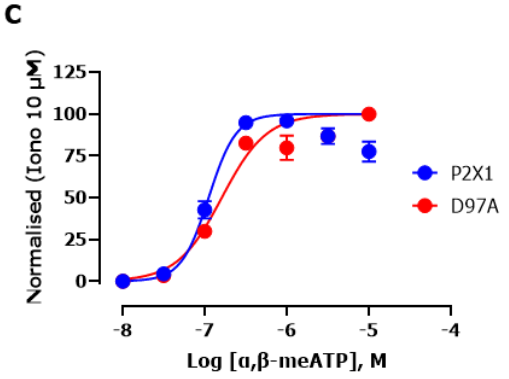


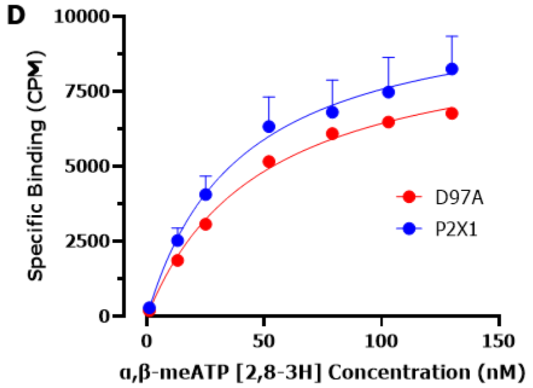


E

| | |
|---|---|
| hP2X1 | D97 |
| hP2X2 | G105 |
| hP2X3 | T87 |
| hP2X4 | E96 |
| hP2X5 | E98 |
| hP2X6 | E107 |
| hP2X7 | - |

**Supplementary Figure 10**

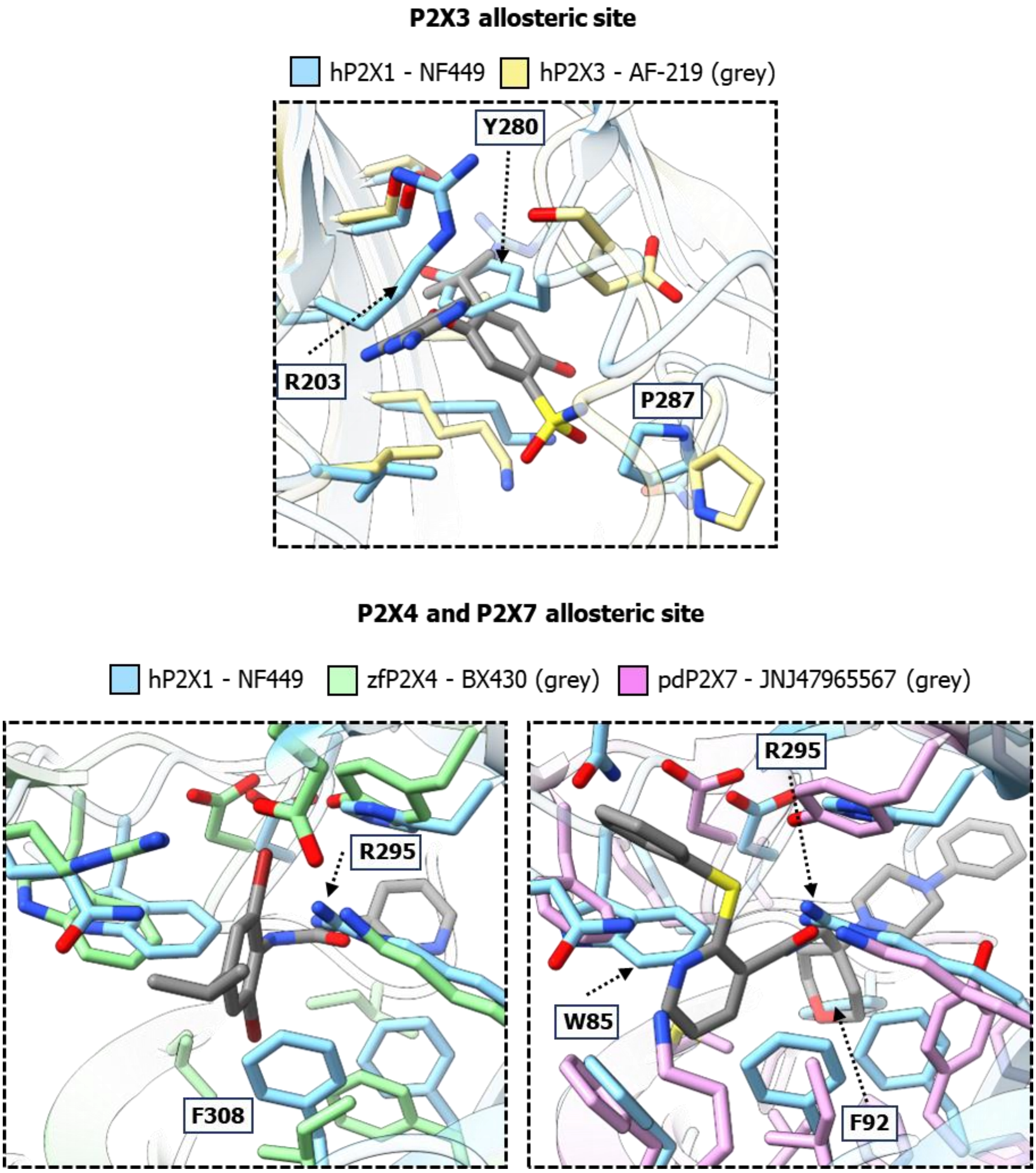

**Supplementary Table 1**

| | ATP-bound P2X1 receptor (9B73) | NF449-bound P2X1 receptor (9B95) |
|---|---|---|
| **Data Collection** | | |
| Micrographs | 6178 + 4578 | 5837 |
| Electron dose ($e^-/A^2$) | 60 | 60 |
| Voltage (kV) | 300 | 300 |
| Pixel size (Å) | 0.82 | 0.82 |
| Defocus range (μm) | 0.5 – 1.5 | 0.5-1.5 |
| Symmetry imposed | C3 | C3 |
| Particles (final map) | 481,309 | 41,932 |
| Resolution (0.143 FSC) (Å) | 1.96 | 2.61 |
| B factor ($Å^{-2}$) | 60.7 | 72.9 |
| **Model Validation** | | |
| Clashscore, all atoms: | 6.02 | 7.18 |
| Poor rotamers | 3 (0.4%) | 0 (0%) |
| Favored rotamers | 837 (99.3%) | 816 (98.6%) |
| Ramachandran outliers | 0 (0%) | 0 (0.00%) |
| Ramachandran favored | 972 (100%) | 945 (98.8%) |
| Rama distribution Z-score | 0.18 ± 0.28 | -1.00 ± 0.28 |
| MolProbity score^ | 1.33 | 1.39 |
| Cβ deviations >0.25Å | 0 (0.00%) | 0 (0%) |
| Bad bonds: | 0 / 8157 (0%) | 0 / 7845 (0%) |
| Bad angles: | 0 / 11088 (0%) | 0 / 10641 (0%) |
| CaBLAM outliers | 6 (0.6%) | 15 (1.6%) |
| CA Geometry outliers | 3 (0.3%) | 6 (0.6%) |
| Cis Prolines: | 6 / 45 (13.3%) | 6 / 45 (13.3%) |
| Chiral volume outliers | 0 / 1242 | 0 / 1179 |
| Waters with clashes | 3 / 66 (4.5%) | 0 / 18 (0%) |

**Supplementary Table 2**

| P2X1 Residue | Mg-ATP | Interaction | Distance |
|---|---|---|---|
| **Lys 68** | α phosphate | Salt bridge | 2.7 |
| | β phosphate | Salt bridge | 3.1 |
| | γ phosphate | Salt bridge | 2.8 |
| **Lys 68 (backbone carbonyl)** | Adenine ring | Hydrogen bond | 2.8 |
| **Lys 70** | γ phosphate | Salt bridge | 2.7 |
| | γ phosphate | Salt bridge | 3.3 |
| **Lys 70** | Adenine ring | Pi-cation bond | 4.8 |
| **Glu 122** | $Mg^{2+}$ ($H_2O$) | Hydrogen bond | 2.7 |
| | $Mg^{2+}$ ($H_2O$) | Hydrogen bond | 2.7 |
| **Arg 139** | Adenine Ring ($H_2O$) | Hydrogen bond | 2.9 |
| **Lys 140** | Ribose sugar | Hydrogen bond | 3.1 |
| **Asp 170** | $Mg^{2+}$ | Ionic bond | 2.0 |
| **Thr 186** | Adenine ring | Hydrogen bond | 2.8 |
| **Thr 186 (backbone carbonyl)** | Adenine ring | Hydrogen bond | 2.7 |
| **Phe 188** | Adenine ring | Pi-Pi stacking | 4.8 |
| **Met 214** | Ribose sugar | Hydrophobic Interaction | 4.0 |
| **Val 229** | Adenine ring | Hydrophobic Interaction | 3.6-3.8 |
| **Ser 286** | α phosphate | Hydrogen bond | 2.3* |
| **Asn 290** | β phosphate | Hydrogen bond | 2.8 |
| **Arg 292** | γ phosphate | Salt bridge | 2.5* |
| | γ phosphate | Salt bridge | 2.7 |
| **Lys 309** | γ phosphate | Salt bridge | 2.8 |
| | β phosphate | Salt bridge | 2.9 |
| **$Mg^{2+}$ (ATP)** | 4x $H_2O$ | Ionic Interaction | 2.0 |
| | Asp 170 | Ionic Interaction | 2.0 |
| | γ phosphate | Ionic Interaction | 2.0 |

**Supplementary Table 3**

| P2X1 Residue | NF449 | Interaction | Distance |
|---|---|---|---|
| **Lys 68** | Sulfonic acid | Salt bridge | 2.8 |
| **Lys 70** | Sulfonic acid | Salt bridge | 3.3 |
| **Leu 72 (backbone amine)** | Sulfonic acid | Hydrogen bond | 2.6* |
| | Sulfonic acid | Hydrogen bond | 2.8 |
| **Leu 72 (backbone carbonyl)** | Sulfonic acid | Hydrogen bond | 2.9 |
| **Lys 136** | Sulfonic acid | Salt bridge | 3.2 |
| **Arg 139** | Benzene ring | Pi-cation bond | 3.9 |
| **Lys 140 (backbone amine)** | Sulfonic acid | Hydrogen bond | 3.2 |
| | Sulfonic acid | Hydrogen bond | 2.8 |
| **Thr 186** | Benzene ring, Amide | Hydrophobic Interaction | 3.3-4.0 |
| **Phe 188** | Benzene ring | Pi-Pi stacking | 4.5 |
| **Val 209** | Benzene ring | Hydrophobic Interaction | 3.5-3.8 |
| **Met 214** | Sulfonic acid | Hydrogen bond | 3.3* |
| **Lys 215** | Sulfonic acid | Salt bridge | 3.5 |
| **Cys 217 (backbone amine)** | Urea | Hydrogen bond | 3.1 |
| **Cys 217** | Benzene ring | Hydrophobic Interaction | 3.9 |
| **Leu 218** | Benzene ring | Hydrophobic Interaction | 3.7-3.9 |
| **Pro 228** | Benzene ring | Hydrophobic Interaction | 3.5-3.8 |
| **Val 229** | Benzene ring, Urea | Hydrophobic Interaction | 3.5-3.8 |
| **Glu 282** | N/A | N/A | N/A |
| **Asn 290** | Sulfonic acid | Hydrogen bond | 3.6* |
| **Arg 292** | Sulfonic acid | Salt bridge | 2.8 |
| | Sulfonic acid | Salt bridge | 3.1 |
